# Dopamine biosynthesis in plants

**DOI:** 10.64898/2025.12.10.693491

**Authors:** Yezhang Ding, Yi Zhai, Tomáš Brůna, Sharon Greenblum, Li Lei, Leo A. Baumgart, Peng Wang, Vlastimil Novak, Mingqin Shao, Scott J. Lee, Samuel P. Hazen, Suzanne M. Kosina, John P. Vogel, Trent R. Northen

## Abstract

Dopamine, an essential human neurotransmitter, also plays important roles in plant-microbe interactions and abiotic stress tolerance. While dopamine biosynthesis is well-characterized in mammals, it remains largely unclear in plants. Through metabolite-based genome-wide association studies, we identified two putative polyphenol oxidase genes for dopamine biosynthesis in *Brachypodium distachyon*, located in the same chromosomal region as two tyrosine decarboxylase genes. Biochemical and knockout mutant analyses demonstrated that dopamine biosynthesis proceeds primarily via tyramine as an intermediate and contributes to plant tissue browning. Homology searches and functional assays indicate that this dopamine biosynthetic pathway is conserved across diverse plant species. This discovery provides valuable insights into plant dopamine biosynthesis with potential applications in reducing food browning, improving stress resilience, and bioengineering valuable compounds.

## Introduction

Dopamine is a critical neurotransmitter in humans, often termed the “feel-good” hormone due to its role in generating sensations of pleasure, satisfaction, and motivation (Mirenowicz and Schultz, 1996). In the brain, dopamine can spontaneously form neuromelanin through melanogenesis, which is a brown-to-black pigment (Hedges et al., 2020; Krainc et al., 2023). Interestingly, dopamine is also synthesized by a wide range of plant species (Gomes et al., 2024; Liu et al., 2020), where it contributes to biotic and abiotic stress tolerance (Ahmad et al., 2021; Liu et al., 2020). Recent studies have shown that dopamine is exuded from roots of the model grass *Brachypodium distachyon* (Novak et al., 2024), and can modulate soil microbial community structure (Ding et al., 2024). Moreover, dopamine is a known precursor to various pharmaceutically valuable metabolites, including phenethylisoquinoline alkaloids, benzylisoquinoline alkaloids, catecholamines, and phenylethylamines (Xu et al., 2020).

In mammals, dopamine biosynthesis involves the hydroxylation of tyrosine to form L-3,4-dihydroxyphenylalanine (L-DOPA), which is subsequently decarboxylated by DOPA decarboxylase (DODC) to produce dopamine (Bromek et al., 2010; Meiser et al., 2013). A minor pathway also exists in mammals, where tyrosine decarboxylases (TyDCs) first decarboxylate tyrosine to produce tyramine, which is subsequently hydroxylated by cytochrome P450 enzymes to form dopamine (Bromek et al., 2010). Similar dopamine biosynthetic routes have been proposed to exist in plants (Facchini and De Luca, 1995; Facchini et al., 2000; Liu et al., 2020) (Supplementary Fig. 1), yet the enzymes involved in planta remain largely unclear (Xu et al., 2020).

Enzymatic browning is a biological process that significantly impacts food quality, particularly fruits and vegetables, reducing shelf-life and thereby leading to food waste. The process is initiated by polyphenol oxidases (PPOs), which catalyze the oxidation of phenolic compounds, including dopamine. The resulting products undergo further non-enzymatic reactions, leading to the polymerization of these compounds into dark pigments known as melanin, often seen on cut bananas, apples, and other produce (Sui et al., 2023; Tilley et al., 2023). Food enzymatic browning can be mitigated using traditional methods, such as lowering pH, reducing oxygen exposure, or applying chemical inhibitors, as well as advanced genome-editing techniques to knock down or knockout *PPO* genes (Sui et al., 2023; Tilley et al., 2023). Nonetheless, understanding of the genetic and metabolic basis of browning remains critical for developing next-generation crops with improved resistance to discoloration and enhanced postharvest quality.

Here, we present the identification of a plant-specific dopamine biosynthetic pathway in *B. distachyon* using a combination of metabolite-based genome-wide association studies (mGWAS), gene expression analysis, enzyme assays, and knockout mutant characterization. Our results reveal that unlike in mammals, dopamine biosynthesis in the model grass plant *B. distachyon* proceeds primarily via tyramine as an intermediate, where TyDCs convert tyrosine into tyramine, which is then converted into dopamine by PPOs. Evidence from homologous searches and functional assays suggests that the dopamine biosynthetic pathway is highly conserved across a wide range of plant species. Notably, dopamine-deficient mutants, including *Bdtydc1Bdtydc2*, *Bdppo1*, and *Bdppo1Bdppo3*, exhibited a lack of dark brown coloration in root tissues after senescence or damage. Additionally, transient expression of the dopamine biosynthetic pathway in *Nicotiana benthamiana* recapitulated the browning phenotype. Together, these findings suggest that the dopamine pathway can be a major component of tissue browning in plants. The elucidation of the plant dopamine biosynthetic pathway offers new opportunities for development of plants with reduced browning as well as production of valuable dopamine-derived compounds among other potential benefits.

## Results

### Identification of candidate genes in dopamine biosynthesis

Given the importance of dopamine in plants, we sought to identify the gene(s) responsible for its biosynthesis. A *B. distachyon* diversity panel, consisting of 100 natural accessions (Supplementary Table 1), was grown in a hydroponic system, and the roots and shoots of 4-week-old plants were harvested for metabolite analysis using LC-MS/MS. In addition to dopamine (Ding et al., 2024), we measured high levels of tyramine and only detected trace levels of L-DOPA in *B. distachyon* (Supplementary Figs. 2 and 3).

To identify the gene(s) for dopamine biosynthesis, we performed metabolite-based genome-wide association studies (mGWAS). When dopamine levels in roots or shoots were directly used as mapping traits, no significant single-nucleotide polymorphisms (SNPs) were detected (Supplementary Fig. 4A and 4B). Given that using the ratio of putative precursor and product metabolite concentrations as mapping traits can reduce biological variability and enhance statistical power in mGWAS (Wu et al., 2023), we instead used the dopamine-to-tyramine ratio as the trait. Following this approach, we identified two overlapping association peaks with significant SNPs on chromosome 2 (Fig. 1A and 1B; Supplementary Fig. 4C and 4D). Within the genetic mapping interval associated with the dopamine-to-tyramine ratio in roots, an examination of the Bd21-3 genome revealed four genes (Fig. 1C). Notably, one of them, *BdiBd21-3.2G0666400*, encodes a putative polyphenol oxidase (PPO) and shows dominant expression in roots (Supplementary Fig. 5; Supplementary Tables 2 and 3), suggesting that *BdiBd21-3.2G0666400* is responsible for dopamine biosynthesis in roots. Interestingly, in the genetic mapping interval associated with the dopamine-to-tyramine ratio in shoots, three putative PPO-encoding genes were identified, *BdiBd21-3.2G0666400, BdiBd21-3.2G0667800,* and *BdiBd21-3.2G0668300* (Supplementary Fig. 4D; Supplementary Table 4). Sequence comparison revealed that the predicted protein encoded by *BdiBd21-3.2G0666400* shares 52.1% and 78.9% sequence identity with the predicted proteins encoded by *BdiBd21-3.2G0667800* and *BdiBd21-3.2G0668300*, respectively (Supplementary Fig. 6). The three putative PPO-encoding genes were, therefore, named *BdPPO1, BdPPO2*, and *BdPPO3* according to their physical order on the chromosome. Gene expression analysis showed that *BdPPO1* and *BdPPO2* were predominantly expressed in the roots of both 2-week-old and 4-week-old *B. distachyon* Bd21-3 plants, whereas *BdPPO3* was mainly expressed in the shoots of both 2-week-old and 4-week-old plants (Fig. 1D).

**Fig. 1.**
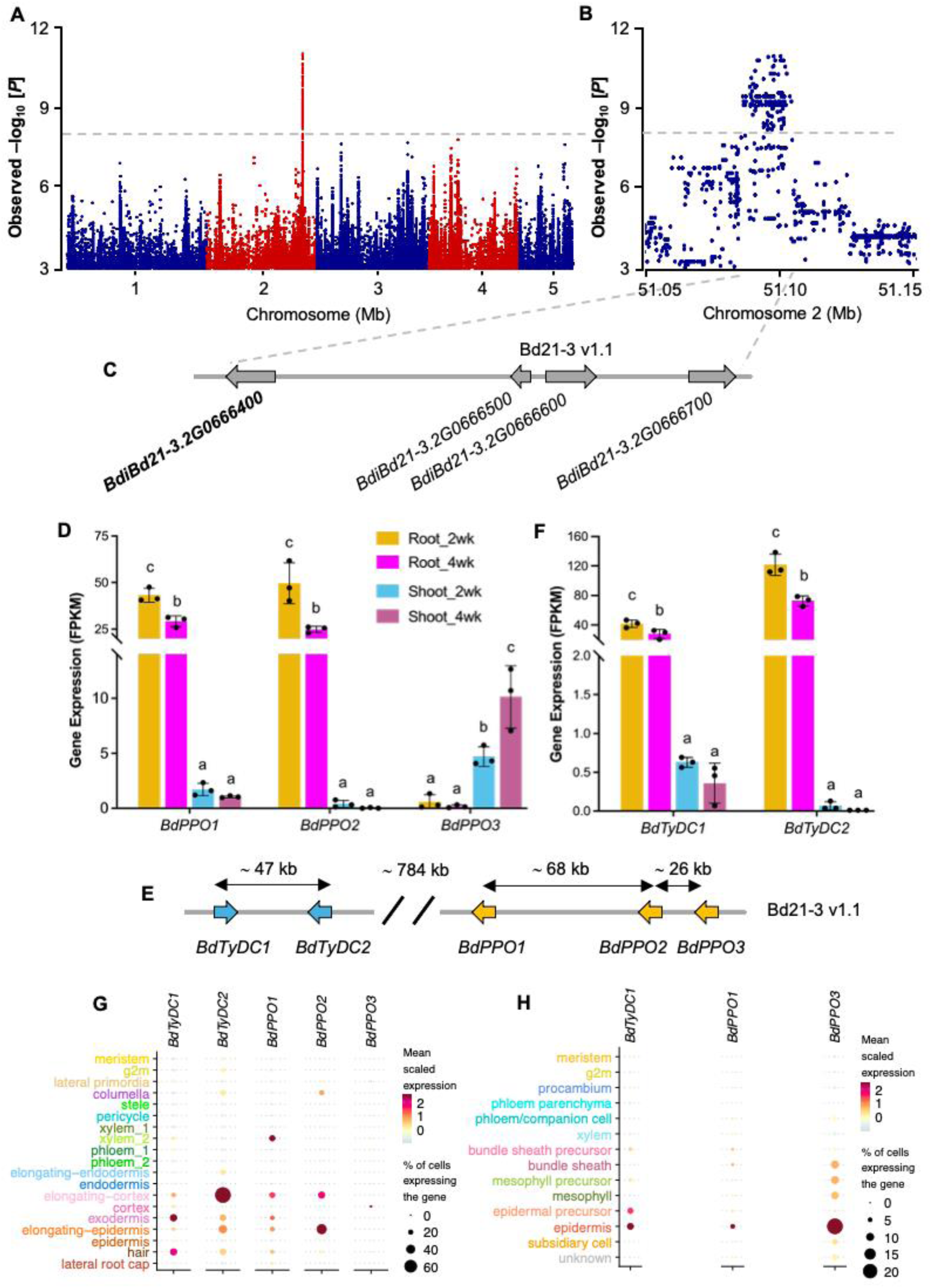
Genetic mapping of candidate dopamine biosynthetic genes in *B. distachyon*. (**A)** Manhattan plot of a GWAS using the dopamine-to-tyramine ratio in roots as the mapping trait. The y-axis shows negative log10-transformed *P*-values from the general linear model. The dashed line represents the 5% Bonferroni-corrected significance threshold. **(B)** A regional Manhattan plot representing a ‘zoomed-in’ view of the signal between 51.05 Mb and 51.15 Mb on chromosome 2. **(C)** Four genes within the mapping region containing SNPs with negative log10-transformed *P*-values exceeding the significance threshold. **(D)** The transcript abundance of three putative *BdPPOs* from RNA-seq analyses of 2-week-old and 4-week-old roots (Root_2wk and Root_4wk) and shoots (Shoot_2wk and Shoot_4wk). **(E)** Three putative *BdPPO*s are co-localized with two *BdTyDC*s on Chromosome 2. **(F)** The transcript abundance of *BdTyDC1 and BdTyDC2* from RNA-seq analyses of 2-week-old and 4-week-old roots (Root_2 and Root_4) and shoots (Shoot_2 and Shoot_4). Gene expression is given as fragments per kilobase of transcript per million mapped reads (FPKM). Error bars represent mean ± SD (*n* = 3 independent samples). For each gene, different letters (a-c) indicate statistically significant differences between samples (one-way ANOVA followed by Tukey’s post hoc test; *P* < 0.05). **(G-H)**, snRNA-seq results showing the candidate genes are differentially expressed in different cell types. Dotplots of normalized, scaled expression of candidate dopamine biosynthetic genes in each annotated cell type of roots (**G**) and shoots (**H**) (*n* = 2 independent samples).

Previous studies revealed that plant TyDCs are involved in dopamine biosynthesis (Facchini and De Luca, 1995; Facchini et al., 2000; Wang et al., 2021). Notably, we found that two known TyDC-encoding genes, *BdiBd21-3.2G0653800* and *BdiBd21-3.2G0654700* (Noda et al., 2015), are located within the same chromosomal region, approximately 829.0 kb and 783.6 kb upstream of *BdPPO1*, respectively (Fig. 1E), suggesting that these physically linked genes may represent a putative dopamine biosynthetic gene cluster in *B. distachyon*. The predicted proteins encoded by the two *TyDC* genes share 97.9% sequence identity. We designated *BdiBd21-3.2G0653800* as *BdTyDC1* and *BdiBd21-3.2G0654700* as *BdTyDC2*. Gene expression analysis revealed that *BdTyDC1* and *BdTyDC2* were predominantly expressed in the roots of 2-week-old and 4-week-old *B. distachyon* Bd21-3 plants (Fig. 1F).

We next performed single-nuclei RNA-seq (snRNA-seq) on roots and leaves of 8-day-old *B. distachyon* plants to determine whether candidate gene expression was limited to specific cell types and whether *BdPPOs* and *BdTyDCs* were co-expressed. After stringent filtering for high-quality cells, we constructed cell type-resolved gene expression atlases consisting of 9,700 root nuclei and 12,234 leaf nuclei (Supplementary Tables 5 and 6). Leveraging published marker gene sets from maize, *Sorghum bicolor*, and rice (Guillotin et al., 2023; He et al., 2024; Zhang and Sun, 2021), we identified all major cell types in our atlases, including vasculature, ground cells, and epidermal cells of varying subtypes and developmental stages in roots, and vasculature, mesophyll, bundle sheath, and epidermis in leaves (Supplementary Fig. 7A and 7B). Consistent with our bulk RNA-seq results, *BdPPO1* was predominantly expressed in roots, while *BdPPO3* was primarily expressed in leaves (Fig. 1G and 1H). Notably, *BdPPO1* expression was specifically limited to developing cortex and exodermis cells, along with a subset of xylem cells expressing genes involved in secondary xylem differentiation (Fig. 1G; Supplementary Tables 7 and 8). *BdPPO2* was also expressed in roots yet exhibited a distinct expression pattern with highest expression in developing epidermis (Fig. 1G and 1H). In contrast, while *BdPPO3* was weakly expressed in mature root cortex cells, it was strongly expressed in leaf mesophyll and epidermis (Fig. 1G and 1H). Additionally, we observed that *BdTyDC1* and *BdTyDC2* were predominantly expressed in roots, particularly in developing cortex and developing epidermal cells (Fig. 1G and 1H). Interestingly, *BdTyDC1* was also expressed in leaf epidermal cells, while *BdTyDC2* expression was completely absent in leaves (Fig. 1G and 1H). Intriguingly, we found that *BdTyDC1* and *BdTyDC2* are the top two genes co-expressed with *BdPPO1* in roots (Supplementary Table 7), further supporting their involvement in root dopamine biosynthesis. Collectively, these findings reveal that these candidate dopamine biosynthetic genes in *B. distachyon* are spatially expressed not only in different tissues but also in distinct cell types within these tissues.

### Characterization of BdTyDCs in dopamine biosynthesis

BdTyDC1 and BdTyDC2 were previously shown to possess L-tyrosine decarboxylation activity in yeast (Noda et al., 2015). To confirm their enzymatic function in planta, we transiently expressed each gene in *N. benthamiana* leaves using *Agrobacterium*-mediated transformation. Expression of either *BdTyDC1* or *BdTyDC2* led to the accumulation of tyramine (Fig. 2A), consistent with the enzymes functioning as tyrosine decarboxylases (Fig. 2B). To further investigate their roles in dopamine biosynthesis, we generated single and double knockout mutants of *BdTyDC1* and *BdTyDC2* using CRISPR-Cas9 gene editing (Supplementary Fig. 8). Knocking out *BdTyDC1* resulted in a 50% reduction in tyramine levels in roots and a 90% reduction in shoots, along with a corresponding decrease in dopamine levels by 30% in roots and 90% in shoots (Fig. 2C and 2D; Supplementary Fig. 9). In contrast, *BdTyDC2* knockout plants exhibited reductions in root tyramine (70%) and dopamine (50%) levels, but showed no changes in shoot levels (Fig. 2C and 2D; Supplementary Fig. 9). Strikingly, the *Bdtydc1Bdtydc2* double mutants showed a complete loss of both tyramine and dopamine in roots and shoots (Fig. 2C and 2D; Supplementary Fig. 9), indicating that these two enzymes act redundantly in the biosynthesis of tyramine, with *BdTyDC1* playing a dominant role in shoot tissues. Additionally, we observed that the *Bdtydc1Bdtydc2* double mutant plants only accumulated trace levels of L-DOPA in roots, comparable to those observed in wild-type plants (Supplementary Fig. 10), suggesting that BdPPOs are unlikely to catalyze the 3-hydroxylation of tyrosine in *B. distachyon* under native physiological conditions. Taken together, our results support a biosynthetic pathway in *B. distachyon* in which dopamine is primarily synthesized via tyramine as an intermediate.

**Fig. 2.**
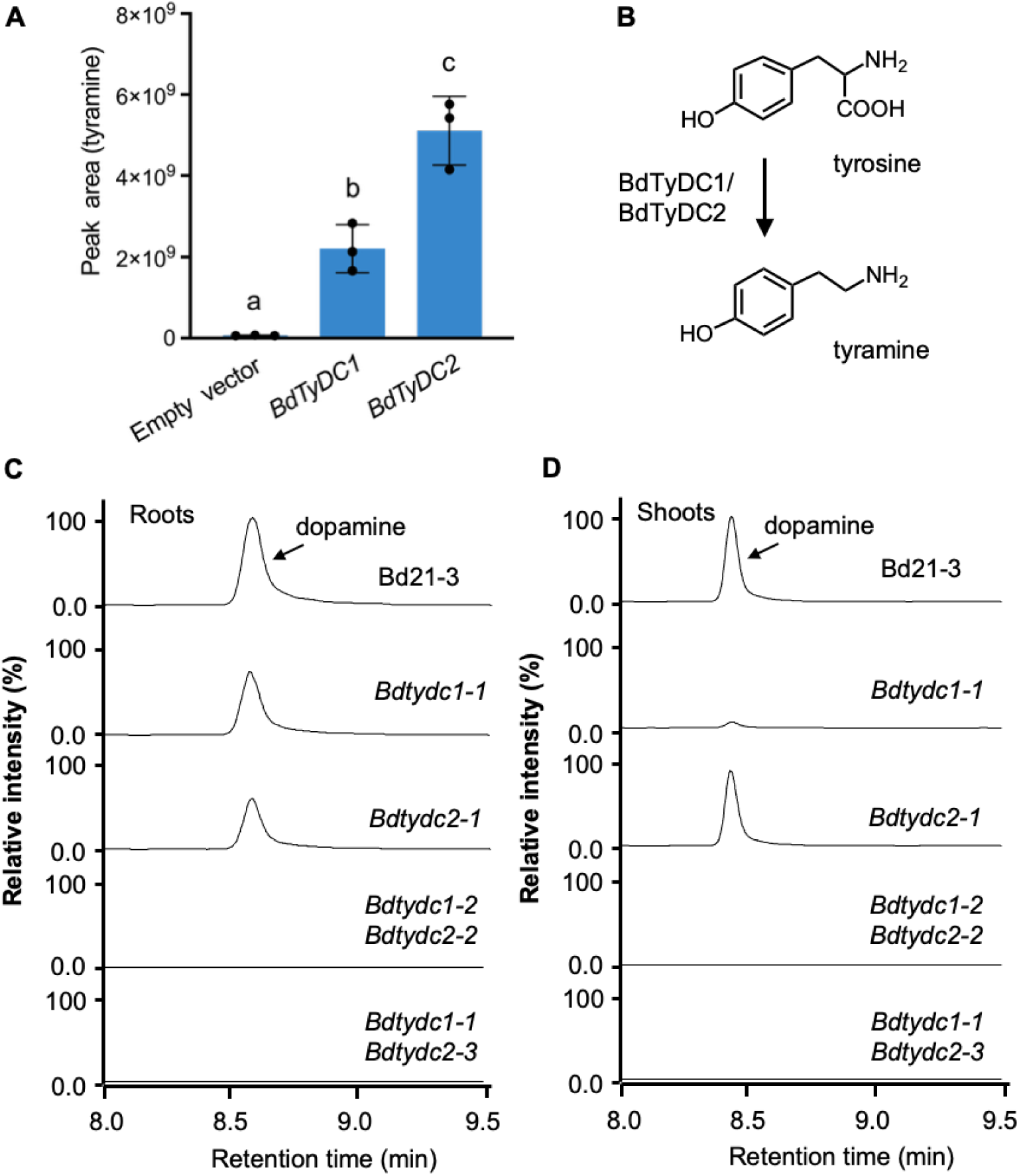
Characterization of BdTyDC1 and BdTyDC2. **(A)** Tyramine levels ([M+H]^+^) in *N. benthamiana* leaves transiently expressing *BdTyDC1*, *BdTyDC2*, or empty vector, as measured by LC-MS/MS. The values indicate means of biological replicates ± standard deviation (SD) (*n* = 3 independent samples). Within the plot, different letters (a-c) represent significant differences (one-way ANOVA followed by Tukey’s test corrections for multiple comparisons; *P* < 0.05). **(B)** Proposed reaction by BdTyDC1 and BdTyDC2 in the biosynthesis of Tyramine. **(C-D)** Comparison of dopamine levels in *Bdtydc* mutant lines. Representative LC-MS extracted ion chromatograms of dopamine ( [M+H]^+^) in roots (**C**) and shoots (**D**) of Bd21-3 (wildtype) and the *Bdtydc* mutants, *Bdtydc1-1*, *Bdtydc2-1*, *Bdtydc1-2Bdtydc2-2,* and *Bdtydc1-1Bdtydc2-3*. Bar plots with biological replicates can be found in Supplementary Fig. 9. Plants were grown in a hydroponic system for 4 weeks, and roots and shoots were harvested for metabolite analysis via LC-MS/MS.

### Characterization of BdPPOs in dopamine biosynthesis

To investigate whether *BdPPO1*, *BdPPO2*, and/or *BdPPO3* possess tyrosinase or tyrosine hydroxylase activity, we transiently expressed each gene individually in *N. benthamiana* leaves using *Agrobacterium*-mediated transformation. Among the three enzymes, only *BdPPO1* expression led to a low but detectable increase in L-DOPA production (Supplementary Fig. 11), suggesting that *BdPPO1* exhibits minor activity for the 3-hydroxylation of tyrosine *in vivo*. To assess their activity in dopamine biosynthesis, we next co-expressed *BdTyDC2* with each of the three *BdPPOs* in *N. benthamiana*. Co-expression of *BdTyDC2* with *BdPPO1*, *BdPPO2*, or *BdPPO3* led to dopamine accumulation and reduced tyramine accumulation (Fig. 3A; Supplementary Fig. 12A, 12B, and 12C), indicating that all three BdPPO enzymes are capable of catalyzing the 3-hydroxylation of tyramine. These results support a model in which BdPPOs hydroxylate tyramine during dopamine biosynthesis (Fig. 3B).

**Fig. 3.**
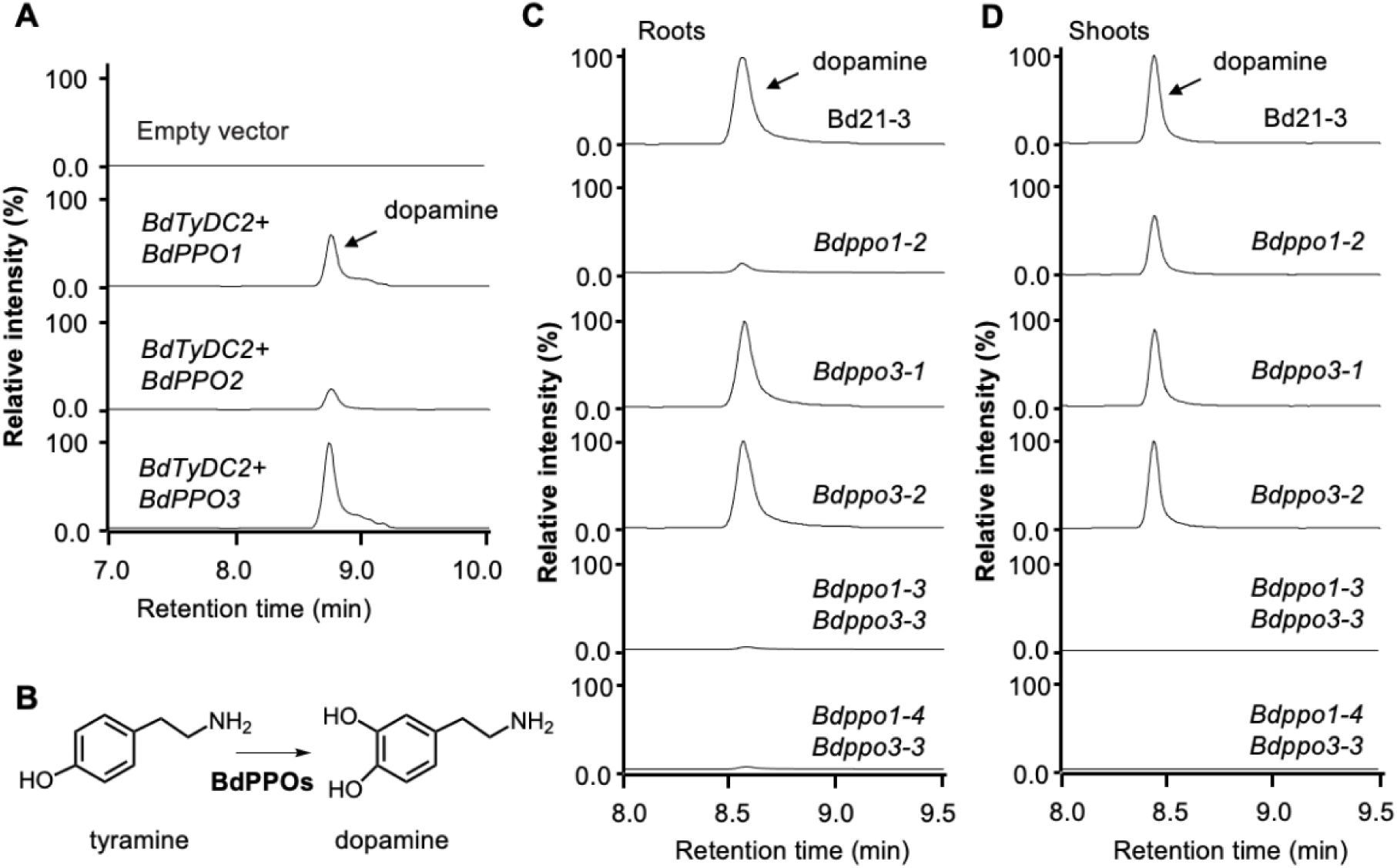
Functional characterization of *BdPPO* genes in dopamine biosynthesis. **(A)** Representative LC-MS extracted ion chromatograms of dopamine ([M+H]^+^) in *N. benthamiana* leaves transiently co-expressing *BdTyDC2* with *BdPPO1, BdPPO2, or BdPPO3*. Bar plots with biological replicates can be found in Supplementary Fig. 12C. **(B)** Proposed enzymatic reaction catalyzed by BdPPOs in the biosynthesis of dopamine from tyramine. **(C-D)** Representative LC-MS extracted ion chromatograms of dopamine ([M+H]^+^) in the roots (**C**) and shoots (**D**) of Bd21-3 (wildtype) and *Bdppo* mutants, including *Bdppo1-2*, *Bdppo3-1*, *Bdppo3-2*, *Bdppo1-3Bdppo3-3, and Bdppo1-4Bdppo3-3*. Bar plots with biological replicates can be found in Supplementary Fig. 9. Plants were grown in a hydroponic system for 4 weeks, and roots and shoots were harvested for metabolite analysis using LC-MS/MS.

To further examine BdPPO1’s role in dopamine biosynthesis, we obtained two *Bdppo1* knockout mutant alleles: *Bdppo1-1* (NaN1435), identified from the publicly available sodium azide *B. distachyon* sequenced mutant population (https://jgi.doe.gov/indexed-collection-of-brachy-mutants/), and *Bdppo1-2*, generated using CRISPR-Cas9 gene editing (Supplementary Figs. 13 and 14). We found that both *Bdppo1* mutant alleles exhibited a severe reduction in root dopamine biosynthesis, while dopamine levels in shoots remained similar to those in wild-type plants (Fig. 3C and 3D; Supplementary Figs. 9B, 9D, 13B, and 13D). In contrast, a knockout mutation in *BdPPO2* did not affect dopamine accumulation in roots or shoots (Supplementary Fig. 15), indicating that dopamine biosynthesis in *B. distachyon* roots is predominantly catalyzed by BdPPO1. Additionally, both *Bdppo1* mutant alleles showed more than a 10-fold increase in root tyramine levels compared to wild-type plants (Supplementary Figs. 9B and 13C), further supporting the hypothesis that dopamine is synthesized via tyramine as an intermediate (Fig. 3B).

To investigate dopamine biosynthesis in shoots, we next generated *Bdppo3* single and *Bdppo1Bdppo3* double knockout mutants using CRISPR-Cas9 gene editing (Supplementary Fig. 14A and 14B). The two *Bdppo3* mutant alleles produced wild-type levels of dopamine in shoots, whereas the *Bdppo1Bdppo3* double mutant plants exhibited a nearly complete deficiency in shoot dopamine accumulation (Fig. 3C and 3D; Supplementary Fig. 9). These results suggest that *BdPPO1* and *BdPPO3* function redundantly in shoot dopamine biosynthesis. Interestingly, dopamine levels in the roots of *Bdppo1Bdppo3* double mutants were significantly lower than those in *Bdppo1* single mutants, indicating that *BdPPO3* also contributes to root dopamine biosynthesis, albeit to a lesser extent (Supplementary Fig. 9). Taken together, our findings demonstrate that both *BdPPO1* and *BdPPO3* are involved in dopamine biosynthesis in *B. distachyon*, with *BdPPO1* playing a predominant role in roots and both enzymes acting redundantly in shoots.

### Dopamine oxidation causes tissue browning in plants

To examine whether the dopamine pathway has an effect on plant growth, we grew these mutant plants in a hydroponic system. We did not observe any growth defects in the biomass of the roots and shoots of the 4-week-old dopamine pathway mutant plants compared with the wild type (Supplementary Fig. 16A and 16B). Interestingly, we observed that the root tissues of the dopamine-deficient plants, including *Bdtydc1Bdtydc2*, *Bdppo1*, and *Bdppo1Bdppo3*, did not turn dark brown after senescence or damage (Fig. 4A and 4B; Supplementary Fig. 17). Moreover, we found that the root extracts of these dopamine-deficient plants did not turn brown after exposure to air for up to a week (Fig. 4C; Supplementary Fig. 18). Previous studies revealed that plant PPOs are responsible for the enzymatic browning of foods, particularly fruits and vegetables, where PPOs catalyze the oxidation of *o*-diphenols to *o*-quinones that further polymerize through non-enzymatic reactions to form brown pigments (Sui et al., 2023; Tilley et al., 2023). Since the browning phenotypes were not observed in the *Bdtydc1Bdtydc2* double mutant plants, and dopamine is known to undergo autoxidation at physiological pH to form brown polymeric products (Umek et al., 2018), we hypothesize that the dopamine pathway causes plant tissue browning. To test our hypothesis, we transiently expressed the dopamine pathway in the leaves of *N. benthamiana*. The extracts from leaves expressing the full dopamine pathway became brown, while those from leaves individually expressing either *BdTyDCs* or *BdPPOs* did not (Fig. 4D). Additionally, we found that the extracts from the leaves of *N. benthamiana* expressing the empty vector developed similar brown coloration after dopamine was added (Fig. 4D). Together, these results strongly suggest that dopamine is an important contributor to the observed tissue browning in plants.

**Fig. 4.**
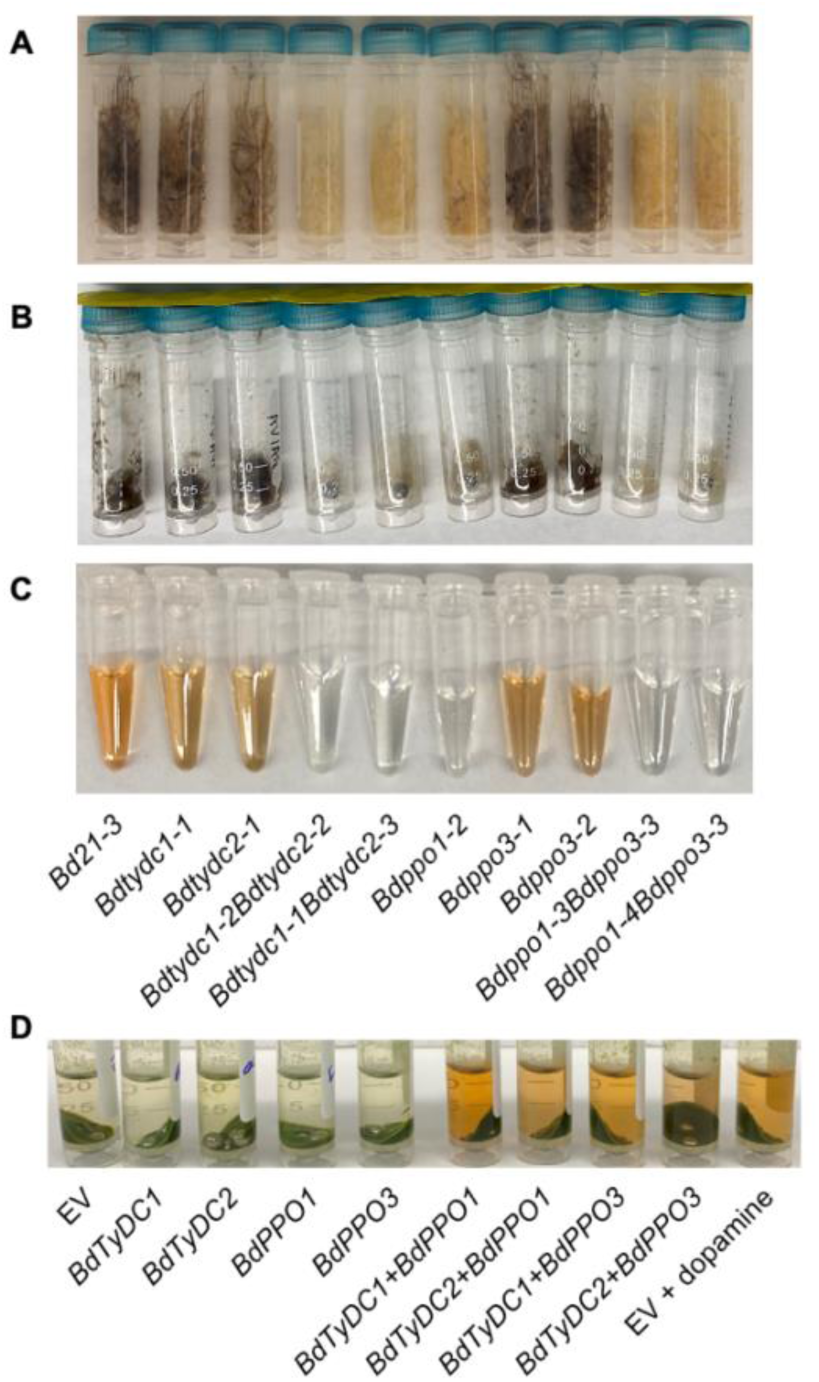
Dopamine oxidation causes tissue browning in plants. **(A)** Root coloration of eight-week-old Bd21-3 (wildtype) and dopamine pathway mutant plants grown on ½ MS plates. **(B)** The coloration of damaged roots (after 2 hours of air exposure) from three-week-old Bd21-3 (wildtype) and dopamine pathway mutant plants grown on ½ MS plates. **(C)** Coloration of root extracts (after 2 hours of air exposure) from three-week-old Bd21-3 and dopamine pathway mutant plants grown on ½ MS plates. The dopamine pathway mutants include *Bdtydc1-1*, *Bdtydc2-1*, two alleles of *Bdtydc1Bdtydc2*, *Bdppo1-2*, two alleles of *Bdppo3*, and two alleles of *Bdppo1Bdppo3.* **(D)** Coloration of leaf extracts (after 2 hours of air exposure) from *N. benthanmiana* transiently expressing *BdTyDC1*, *BdTyDC2*, *BdPPO1*, *BdPPO3*, *BdTyDC1* + *BdPPO1*, *BdTyDC1* + *BdPPO3*, *BdTyDC2* + *BdPPO1*, and *BdTyDC2* + *BdPPO3*. An empty vector (EV) was used for the control.

### The conservation of the dopamine biosynthetic pathway in plants

To investigate the conservation of the dopamine biosynthetic pathway in plants, we searched for *TyDC* and *PPO* gene homologs across flowering plant species using genome sequences available on Phytozome (Goodstein et al., 2012). We found that all examined plant species have putative *TyDC* genes, whereas most plant species, except those in the Brassicaceae family, contain putative *PPO* genes (Fig. 5A). Interestingly, 79 out of 121 (65%) plant species encoding both putative TyDCs and PPOs have at least one pair of them located within 10 Mb on the same chromosome (Fig. 5A). Notably, several plant species prone to browning issues and known for producing high levels of dopamine in their fruits, including *Musa acuminata* (banana) and *Persea americana* (avocado) (Briguglio et al., 2018), were found to possess the dopamine biosynthetic gene homologs localized in the same chromosomal region (less than 10 Mb apart) (Fig. 5A). The enzymatic activity of the TyDC family has been characterized in plants (Facchini and De Luca, 1995; Facchini et al., 2000; Wang et al., 2021); however, the activity of the PPO family remains underexplored in dopamine biosynthesis. To investigate the conserved function of PPOs in dopamine biosynthesis, we examined the activity of select PPOs from several plant species, including *Zea mays*, *Oryza sativa*, *S. bicolor*, *Hordeum vulgare*, and *Papaver somniferum*. Our findings revealed that at least one *PPO* from each select species has the capability to catalyze dopamine production when transiently co-expressed with *BdTyDC2* in the leaves of *N. benthamiana* (Fig. 5B and 5C), indicating a conservation of the dopamine biosynthetic pathway in plants.

**Fig. 5.**
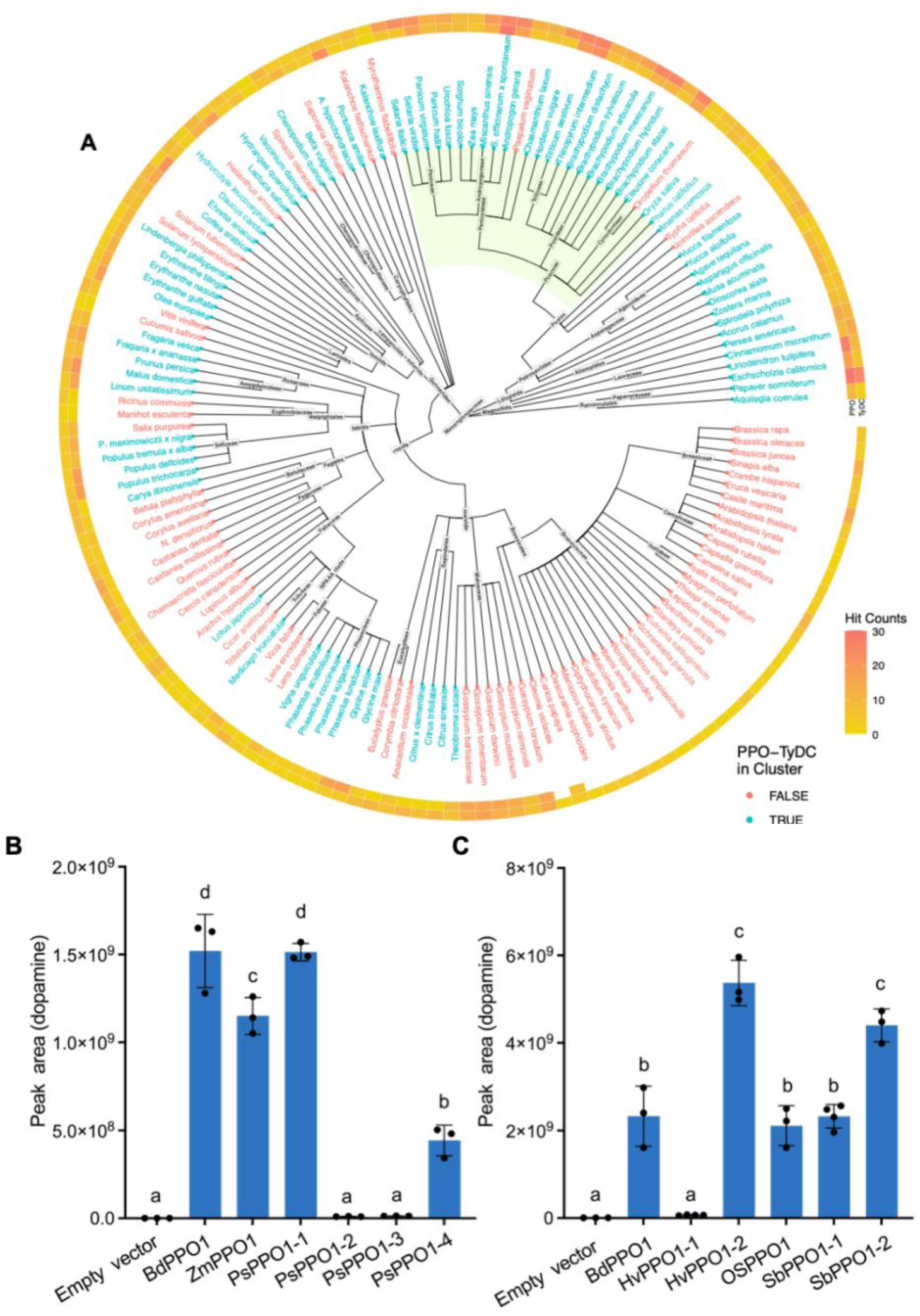
Conservation of the dopamine biosynthetic pathway in plants. **(A)** Numbers of homologous genes in the *TyDC* and *PPO* gene families across flowering plants with sequenced genomes available on Phytozome, as well as *Papaver somniferum*, *Musa acuminata*, and *Persea americana*. Species highlighted in blue possess at least one TyDC-PPO gene pair located within 10 Mb on the same chromosome. **(B-C)** Functional characterization of selected PPOs in dopamine production. *PPOs* from *Zea mays* (*Zm*), *Papaver somniferum* (*Ps*), *Hordeum vulgare* (*Hv*), *Oryza sativa* (*Os*), and *Sorghum bicolor* (*Sb*) were transiently coexpressed with *BdTyDC2* in *N. benthamiana* leaves for five days. Dopamine levels ([M+H]^+^) were analyzed using LC-MS analysis. Error bars represent mean ± SD (*n* = 3 or 4 independent samples). Within the plot, different letters (a-d) represent significant differences (one-way ANOVA followed by Tukey’s test corrections for multiple comparisons; *P* < 0.05).

## Discussion

We report the identification of the genes responsible for dopamine biosynthesis in *B. distachyon*, revealing a plant-specific pathway that diverges from the canonical mammalian pathway. These dopamine biosynthetic genes are located within the same chromosomal region but exhibit distinct tissue-specific expression patterns. Biochemical and genetic evidence demonstrates that, unlike in mammals, dopamine biosynthesis in plants proceeds primarily via tyramine as an intermediate. Homology searches and functional assays suggest that this pathway is conserved across diverse plant species.

PPO-mediated enzymatic browning significantly compromises the quality and shelf life of food, particularly fruits and vegetables (Sui et al., 2023). In this study, we observed that root tissues of dopamine-deficient mutants, including *Bdtydc1Bdtydc2*, *Bdppo1*, and *Bdppo1Bdppo3*, did not develop dark brown coloration upon senescence or damage. Additionally, transient expression of the dopamine pathway (both *TyDC* and *PPO*) resulted in browning in leaf extracts of *N. benthamiana*. These findings demonstrate that the dopamine pathway plays a key role in plant tissue browning. Our results also implicate dopamine oxidation as a likely contributor to browning in widely consumed crops, such as bananas and potatoes, which harbor putative homologs of the identified biosynthetic genes (Fig. 5A) and are known to accumulate high levels of dopamine (Liu et al., 2020). However, this hypothesis requires further experimental validation in these species.

Genome editing techniques, such as RNA interference (RNAi) and CRISPR/Cas9, have been employed to prevent food browning by targeting PPO-encoding genes in various plant species (Tilley et al., 2023). However, our study shows that disrupting PPOs leads to hyperaccumulation of tyramine, a compound with known hypertensive effects in humans (Rafehi et al., 2019). Our results offer a new perspective for bioengineering efforts to reduce browning in plants by targeting alternative gene families, such as those encoding TyDCs, to minimize tyramine accumulation while still reducing browning. Notably, dopamine biosynthetic genes exhibit strong tissue-specific expression patterns, suggesting that their regulation is spatially controlled. This opens the possibility of engineering dopamine pathway activity in targeted tissues, rather than throughout the entire plant. Such tissue-specific manipulation would allow for the reduction of undesirable browning in edible organs (e.g., tubers or fruits), while preserving dopamine-associated defense functions in other plant parts. This strategy provides a refined approach to balance food quality and plant resilience. Beyond its implications for food quality, this work also highlights broader opportunities for engineering the dopamine pathway to improve plant stress resilience. Given dopamine’s potent antioxidant properties, enhancing its biosynthesis in specific tissues could help mitigate oxidative damage caused by reactive oxygen species (ROS) that accumulate under abiotic stress conditions such as drought and salinity (Liu et al., 2020). Moreover, the elucidation of this dopamine biosynthetic pathway enables the use of synthetic biology to engineer both microbial and plant platforms for the sustainable production of dopamine-derived metabolites, including high-value compounds like morphinan alkaloids(Roberts et al., 1987; Vavricka et al., 2022).

## Methods

### Plant materials and plant growth

A *B. distachyon* diversity panel, consisting of 100 natural genetic lines (Supplementary Table 1), was used in this study. *B. distachyon* seeds were sterilized in 70% (v/v) ethanol for 30 s and 5 min in 6% (w/v) sodium hypochlorite, followed by five washes with sterile water. The seeds were subsequently stratified on 1.5% (w/v) phytoagar plates containing 0.5 × Murashige & Skoog (0.5 × MS, MSP01, Caisson Laboratories, USA) and placed in the dark at 4°C for three days. The seeds were then germinated in a growth chamber with a temperature of 22°C, photosynthetic photon flux density (PPFD) at 150 μmol m^-2^s^-1^, and a 16 h light/8 h dark photoperiod for two days.

The culture vessels (PTL-100™, PhytoTech Labs) were rinsed five times with MilliQ water and autoclaved. Two seedlings of each *B. distachyon* line were placed onto a floating plastic holder and then transferred to each vessel with 40 mL of 0.5 × MS. Plants were grown in a 16 h light/8 h dark regime at 22 °C and 50% relative humidity with 150 μmol m^-2^s^-1^ PPFD for four weeks. The culture vessels without plants, filled with the growth medium, were incubated in the same conditions as the controls. At week four, root and shoot tissues were harvested and stored at –80°C for further analysis. The sterility of the hydroponics setup was examined by plating 50 μL medium on Luria Broth (LB, Sigma-Aldrich) solid plates, followed by seven-day incubation at 30°C.

For snRNA-seq experiments, *B. distachyon* Bd21 seeds were surface sterilized by soaking in 75% ethanol for 1 minute and 50% bleach for 8 minutes, 25% bleach with 0.2% Triton X-100 (Sigma-Aldrich 93443) for 8 minutes, and then rinsed with sterile water 5 times. After sterilization, seeds were planted vertically on agar plates containing 3 mM Ca(NO_3_)_2_, 1.5 mM MgSO_4_, 1.25 mM NH_4_H_2_PO_4_, 1 mM KCl, and 1× micronutrients (MS Salts 100×, MSP18-10LT), in 0.8% agar with pH 5.7. Plants were grown in a Percival chamber under conditions of 22°C with 158 μmol m^-2^s^-1^ PPFD on a cycle of 16 h light/8 h dark.

### Metabolite analysis and identification with LC-MS/MS

*B. distachyon* root and shoot tissues were harvested from 4-week-old hydroponically grown plants and stored at –80°C for further analysis. Before extraction, tissues (approximately 0.1 g fresh weight) were ground into powder using bead milling in a Biospec Minibeadbeater-96. 500 μL of extraction buffer (70% methanol, methanol: H_2_O, 70:30, v/v) was then added to the tissue powders for metabolite extraction overnight at 4°C. Alternatively, an acidic extraction buffer [95% (70% methanol, methanol:H_2_O_2_, 70:30, v/v) and 5% 1 M hydrochloric acid] was used to minimize dopamine oxidation. The samples were subsequently centrifuged for 10 min at 15,000 g. The supernatants were dried in a speed vac, resuspended with 150 μL pure methanol, and filtered with 0.22-μm polyvinylidene difluoride microcentrifuge filtration devices (Pall Corporation, NY, USA) (10,000g for 5 min at 10°C). Filtrates were used for metabolite analysis. Polar metabolites were separated using hydrophilic interaction chromatography (HILIC) and detected on a Thermo Q Exactive Hybrid Quadrupole-Orbitrap Mass Spectrometer. Briefly, an Infinity Lab Poroshell 120 HILIC-Z, 2.1×150 mm, 2.7 um column equipped on an Agilent 1290 HPLC stack was used for separations. Data were collected using data-dependent MS2 acquisition to select the top two most intense ions not previously fragmented within 7 seconds. Internal and external standards were used for quality control purposes. Method parameters are defined in the Supplementary Table 9. Using the conditions defined above, dopamine, tyramine, and L-DOPA were identified by comparing their *m*/*z*, retention times, and MS/MS spectra (manual inspection) with those of standards (Sigma-Aldrich) using the MetAtlas workflow (https://github.com/biorack/metatlas) (Yao et al., 2015). Quantification curves, linear regressions, and boxplots were generated using R. The RaMS package was used to extract and visualize extracted ion chromatograms (EICs) and MS/MS spectra. EICs and MS/MS precursor ions were extracted using a 5-ppm mass tolerance window, while fragment ions were matched using a 15-ppm window based on the MS/MS spectrum with the highest precursor intensity within the integrated EIC region. Total and extracted ion chromatograms (TICs and EICs) were prepared using the FreeStyle software (FreeStyle™ 1.5, Thermo Fisher Scientific).

### Metabolite-based genome-wide association studies

Previously, we observed genetic variation in dopamine levels in root exudates of ten *B. distachyon* lines (Ding et al., 2024). To map the gene(s) involved in dopamine biosynthesis, a *B. distachyon* diversity panel, consisting of 100 lines, was grown in a hydroponic system for four weeks, and roots and shoots were harvested for metabolite analysis using LC-MS/MS. Metabolite-based GWAS (mGWAS) were performed using the peak area of dopamine (*m*/*z*, 154.086, [M+H]^+^) and the ratios of dopamine (*m*/*z*, 154.086, [M+H]^+^) to tyramine (*m/z*, 138.091, [M+H]^+^) in roots and shoots as mapping traits. All data were log_2_ transformed prior to statistical analysis to improve normality. SNP (single-nucleotide polymorphism) markers were identified from Illumina sequencing reads of *B. distachyon* natural accessions (Gordon et al., 2020; Gordon et al., 2017; Scarlett et al., 2023). Reads were aligned to the Bd21-3 v1.1 reference genome (https://phytozome-next.jgi.doe.gov/info/BdistachyonBd21_3_v1_1) using BWA (v0.7.17)(Li, 2013; Li and Durbin, 2009) and filtered with Picard tools (v2.18) using FixMateInformation and MarkDuplicates (https://broadinstitute.github.io/picard). GATK (v3.6) was used for base-quality score recalibration, indel realignment, and small InDel discovery using standard hard filtering parameters from GATK Best Practices recommendations (McKenna et al., 2010). The variants filtering and parameter settings are documented in: https://github.com/lilei1/Brachy_mutant/blob/master/analysis/Natural_accessions/filtering_variants.md. In total, 4,780,736 SNPs were retained, with <20% missing genotypes and a minor allele frequency >15%. mGWAS were initially conducted using the General Linear Model (GLM) in TASSEL 5.0 (https://tassel.bitbucket.io/), with final analyses performed using the R package GAPIT (Turner, 2018). Manhattan plots were constructed in the R package qqman (v0.1.4) (http://cran.r-project.org/web/packages/qqman) (Turner, 2018).

### RNA isolation and bulk RNA-seq analyses

Root and shoot tissues were harvested from hydroponically grown *B. distachyon* Bd21-3 plants at weeks 2 and 4. Total RNA was isolated with a NucleoSpin^TM^ RNA Plant Kit (MACHEREY–NAGEL™) according to the manufacturer’s protocol. Three biological replicates were performed using different plants. RNA quality was assessed based on RNA integrity number using an Agilent Bioanalyzer.

RNA-seq library construction and sequencing were performed by Novogene. Messenger RNA (mRNA) was isolated from total RNA using poly-T oligo-attached magnetic beads. Following fragmentation, the first strand of cDNA was synthesized using random hexamer primers, and the second strand of cDNA was then synthesized using either dUTP for directional library preparation or dTTP for non-directional library preparation. The libraries were assessed for quantification using Qubit and real-time PCR, and the size distribution was analyzed with a Bioanalyzer. The quantified libraries were pooled and sequenced on Illumina platforms, based on the effective library concentration and desired data yield. Clustering of the index-coded samples was carried out according to the manufacturer’s instructions. Once cluster generation was complete, the libraries were sequenced on an Illumina platform, producing paired-end reads. The raw reads in fastq format were initially processed using Perl scripts. During this step, clean reads were obtained by removing reads containing adapters, reads with poly-N sequences, and low-quality reads from the raw data. Simultaneously, Q20, Q30, and GC content of the clean data were calculated. Paired-end clean reads were then aligned to the reference genome (*B. distachyon* B21-3 v1.1) using Hisat2 (v2.0.5) (Kim et al., 2015). FeatureCounts (v1.5.0-p3) was used to count the reads numbers mapped to each gene (Liao et al., 2014). The FPKM (Fragments Per Kilobase of transcript sequence per Million base pairs) for each gene was then calculated based on the gene’s length and the number of reads mapped to it using HTSeq (v0.6.1) (Anders et al., 2015).

### Single-nuclei RNA-seq and Analyses

Leaf and root samples for snRNA-seq were harvested from 8-day-old plants at the two-leaf stage. For root samples, sections of 2-4 cm in length beginning from the primary root tips were harvested. For leaf samples, the entire leaf including a small section of coleoptile was harvested. Tissue samples were harvested at 2.5 hours after the start of the day cycle, immediately flash-frozen in liquid nitrogen, and then stored at –80°C for approximately 2.5 months before nuclei isolation. Nuclei isolation from tissues was performed as follows:

Buffer 1 (lysis buffer) consisted of 0.275 M sorbitol (Sigma-Aldrich S6021), 0.1% Triton X-100 (Sigma-Aldrich 93443), 0.01 M MgCl_2_ (Ambion AM9530G), 1× protease inhibitor cocktail (Sigma-Aldrich 4693132001), 0.3 U/μL RNase inhibitor (Roche 03335399001), and 1 mM DTT (Teknova D9750).

Buffer 2 (wash and resuspension buffer) consisted of 1× phosphate buffer saline (without Mg and Ca, Corning, 21-040-CM), Spermidine trihydrochloride 0.5mM (S2501-25G), Spermine tetrahydrochloride 0.1mM (S1141-10G), 0.4 U/μL RNase inhibitor, and 1 mM DTT.

All nuclei isolation steps were carried out in a cold room at 4°C. For each sample, 20-30 mg frozen tissue was chopped for 3 minutes using a razor blade on a glass plate with 100-200 μL buffer 1, then transferred to a petri dish with 1.5 mL cold buffer 1 and allowed to rest on ice with gentle shaking for 2 minutes. The samples were then transferred to a 48-well filter plate (25 μm, 4 mL, Agilent, 201003-100), which was pre-wet with 1 mL cold buffer 1 before it was placed on the receiving plate attached to a QIAvac96 (Qiagen). Pressure was maintained below 100 bar during filtration. The filtered nuclei solution was centrifuged at 500 g for 10 minutes at 4°C to pellet the nuclei, after which the supernatant was discarded. The pellet was resuspended by gentle flicking and pipetting, washed with 2 mL cold buffer 2, centrifuged at 500 g for 5 minutes, and the supernatant was discarded, after which the pellet was again resuspended in 100 μL cold buffer 2. Nuclei quality and quantity were assessed by flow cytometry (BD Accuri C6 plus), for which they were stained by propidium iodide (Sigma-Aldrich P4864) added to a final concentration of 50 μg/mL. The nuclei suspension was then diluted to a concentration of 200-500 nuclei/μL using cold buffer 2.

Single nuclei were partitioned and barcoded on a BD Rhapsody system (BD Rhapsody HT Xpress, BD Rhapsody Scanner), using the BD Rhapsody WTA V3 kit and cartridge, targeting 10,000-25,000 *B. distachyon* nuclei for each of the four samples (two replicate root samples, two replicate leaf samples). Note that samples were multiplexed on the same lane of the cartridge with samples from several other plant species, which were resolved independently by mapping to their respective reference genomes after sequencing. Library creation was carried out following the manufacturer’s protocol. Sequencing was performed on an Illumina NovaSeq X plus 25B flow cell, using 2×150 bp paired-end sequencing and targeting an average of at least 50,000 reads (25,000 fragments) per nucleus. The resulting sequences were pre-processed by filtering out ribosomal RNA using BBTools v38.96 bbduk.sh (BBMap) to remove read pairs with 31-mers in read 2 matching common ribosomal 31-mers. Remaining reads were processed with the BD Rhapsody Sequence Analysis Pipeline v2.2.1 (https://bitbucket.org/CRSwDev/cwl/src/master/), supplied with a combined-species reference genome that included the Bd21 v3.2 reference genome (https://phytozome-next.jgi.doe.gov/sorghumpan/info/Bdistachyon_v3_2) for *B. distachyon*. Separately, rRNA-filtered reads were also mapped to the same reference with STARsolo (Kaminow et al., 2021) in Velocyto mode (‘--soloFeatures Velocyto’) in order to determine splice rates for each cell barcode. Additional QC metrics were compiled by running the R package diem (Alvarez et al., 2020) to assign a ‘debris score’ to each cell barcode, and by fitting a model to the proportion of each species contributing UMIs to each cell barcode (Baumgart et al., 2024) in order to identify putative empty wells and estimate ambient RNA levels. UMIs mapping the organellar genomes were quantified and removed from the analysis, and then cell barcodes were then filtered to those with at least 500 UMIs, at least 250 genes, 90% non-organelle UMIs, 5% unspliced UMIs, debris score <=2, and estimated ambient RNA rate <50%. Count matrices for all remaining cell barcodes were loaded into Seurat v5.0.0 (Hao et al., 2024), and SCTransform v2 normalization was performed, followed by an initial clustering using Seurat’s RunPCA (with 5,000 variable genes and 30 PCs), FindNeighbors, and FindClusters (resolution 0.6). Next, DoubletFinder (McGinnis et al., 2019) was run and cells with a doublet score >0.4 were removed. Finally, the two biological replicates from each tissue were merged into final tissue-specific atlases, which were then re-normalized with SCT before performing a final PCA (30 PCs) and clustering (resolution = 1.0). Cell type labels were assigned to each cluster in each atlas by first examining the expression of a small number of FISH-validated Sorghum bicolor, maize, and rice orthologs from previously published studies (Guillotin et al., 2023; Zhang et al., 2021). Additional cell types were identified using larger sets of previously computed marker genes from the same studies as well as scPlantDB (Hao et al., 2024). Final cell type labels were compiled from these two sources, with additional manual annotation based on literature support for data-derived markers when there were discrepancies or no significant matches.

### 5’ RACE cDNA library construction and gene cloning

Total RNA was isolated from roots and shoots of 4-week-old hydroponically grown Bd21-3 plants using a NucleoSpin^®^ RNA Plant Kit (Takara Bio USA) according to the manufacturer’s protocol. Total RNA, approximately 2 μg, was subjected to TURBO DNA-free treatment (Ambion) and used for the construction of a 5′ rapid amplification of cDNA ends (RACE) cDNA library with the SMARTer RACE 5′/3′ Kit (Clontech) in accordance with the manufacturer’s protocol. Genes with full-length open reading frames (ORFs) were amplified using gene-specific oligonucleotides (Supplementary Table 10). For *Agrobacterium*-mediated transient expression in *N. benthamiana*, full-length open reading frames, including *BdTyDC1*, *BdTyDC2*, *BdPPO1*, *BdPPO2,* and *BdPPO3*, were cloned from cDNA libraries into the expression vector pLIFE33 using the Uracil-Specific-Excision-Reaction (USER) method with USER cloning-specific primers (Geu-Flores et al., 2007) (Supplementary Table 10). The homologous genes of *B. distachyon PPO1* from other species, including *ZmPPO1* (*Zea mays*), *PsPPOs* (*PsPPO1, PsPPO2, PsPPO3*, and *PsPPO4*, *Papaver somniferum*), *HvPPOs* (*HvPPO1* and *HvPPO2*, *Hordeum vulgare*), *OsPPO1* (*Oryza sativa*), and *SbPPOs* (*SbPPO1* and *SbPPO2*, *Sorghum bicolor*), were synthesized (Twist Biosciences) and subcloned into pLIFE33. All native and synthetic gene sequences used in this study for enzyme characterization are detailed in Supplementary Table 11.

### Transient expression in *N. benthamiana*

For transient expression in *N. benthamiana*, full-length open reading frames cloned into the pLIFE33 vector were introduced into *Agrobacterium tumefaciens* strain GV3101. The transformed bacterial cells were cultured at 28°C for 24 hours in LB liquid medium (Sigma-Aldrich) supplemented with 50 mg/L kanamycin, 30 mg/L gentamicin, and 50 mg/L rifampicin. The cells were then harvested and resuspended to a final OD_600_ of 0.8 in a 10 mM MES buffer containing 10 mM MgCl_2_. For all samples except the *BdTyDc2* plus *BdPPO2* samples in Supplementary Fig. 12C, equal volumes of the various cultures, along with the silencing suppressor strain P19, were mixed and infiltrated into the newly expanded leaves of 6-week-old *N. benthamiana* plants using a needleless syringe. For the *BdTyDc2* plus *BdPPO2* samples in Supplementary Fig. 12C, two biological replicates contained twice the volume of the *BdPPO*2 culture. Since the replicates with the larger volume of *BdPPO2* were very similar to the replicate with equal volumes, we presented all three replicates in the Figure. Five days after infiltration, the *Agrobacterium*-infiltrated leaves were harvested for metabolite analysis using LC-MS/MS.

### Identification of the sodium-azide-induced *Bdppo1-1* and *Bdppo2* mutants

Seeds of *Bdppo1-1* (NaN1435, containing a premature stop codon, W382*, in *BdPPO1*) and *Bdppo2* (NaN1956, containing a premature stop codon, E487*, in *BdPPO2*) were obtained from the publicly available sodium azide-induced *B. distachyon* sequenced mutant population (https://jgi.doe.gov/indexed-collection-of-brachy-mutants/). The mutations were verified by sequencing PCR products amplified with gene-specific primers (Supplementary Table 12).

### Generation of dopamine pathway mutants using CRISPR/Cas9

Dopamine biosynthetic gene mutants were generated using CRISPR/Cas9 gene editing. Two gRNAs were designed to simultaneously target *BdTyDC1* and *BdTyDC2*, resulting in single or double knockouts (Supplementary Table 13). For the creation of *Bdppo1*, *Bdppo3*, and *Bdppo1Bdppo3* mutants, one gRNA was designed to target both *BdPPO1* and *BdPPO3* and another to target *BdPPO3* (Supplementary Table 13). Oligonucleotides were phosphorylated and annealed using T4 polynucleotide kinase (New England Biolabs), followed by ligation into the CRISPR destination vector JD633 (Addgene) that was linearized by AarI (New England Biolabs) using a Golden Gate Reaction. All constructs were sequence-verified and introduced individually into *A. tumefaciens* strain AGL1 via electroporation. The transformed bacterial cells were cultured at 28°C for 48 hours on LB agar plates (Sigma-Aldrich) supplemented with 50 mg/L kanamycin, 50 mg/L carbenicillin, and 25 mg/L rifampicin.

*B. distachyon* (Bd21-3) transformation was performed following an established protocol (Bragg et al., 2014). Briefly, plants were grown in a walk-in growth chamber under a long-day photoperiod (16 h light/8 h dark) at 22°C during the day and 26°C at night. Immature seeds were isolated from spikes of approximately 7-week-old plants and sterilized using 0.6% (v/v) sodium hypochlorite and 0.1% Triton X-100 for 4 minutes. After sterilization, seeds were washed 5 times with sterile water to remove residual chemicals. Immature embryos were isolated under a stereomicroscope and placed on a callus induction medium (CIM) containing Linsmaier and Skoog basal medium (PhytoTech Labs) and sucrose. Callus induction was carried out in the dark at 28°C for 4 weeks, with calli subcultured twice to promote embryogenesis. Prior to transformation, *Agrobacterium* carrying the CRISPR/Cas9 construct was streaked onto LB plates and incubated overnight at 28°C.

Bacterial cells were then harvested by scrapping off the plates, resuspended in liquid CIM to an OD_600_ of 0.6, and used to inoculate the embryogenic calli. The calli were immersed in the bacterial suspension for 5 minutes, briefly dried on filter paper, and incubated in the dark at 22°C for 3 days. Following co-culture, the calli were transferred to selective CIM plates containing 40 mg/L hygromycin and 300 mg/L Timentin (PhytoTech Labs) and incubated at 28°C for 10 days under a 16 h light/8 h dark photoperiod and 150 μmol m⁻² s⁻¹ PPFD. Selected calli were then transferred to a regeneration medium containing MS basal medium with vitamins, maltose, 40 mg/L hygromycin, and 300 mg/L Timentin. Regeneration was conducted at 28°C under the same light and humidity conditions for 2-4 weeks. Once shoots were fully developed, regenerated plants were transferred to a rooting medium containing MS basal medium with vitamins and sucrose and incubated at 28°C for 2 weeks. Plants with well-developed roots were transferred to soil for further growth.

Leaves from T0 transgenic plants were collected for genotyping using the Phire Plant Direct PCR kit (Thermo Scientific). Mutations in the target gene were confirmed with gene-specific primers. Homozygous mutant plants were identified in the T1 generation by genotyping the target gene. To prevent sustained off-target effects, T1 homozygous mutants were further genotyped to confirm the absence of the Cas9-containing transgene by PCR amplification of the selectable marker gene (hygromycin B phosphotransferase). Details of the primers used for genotyping are provided in Supplementary Table 12. All sequencing was performed using Sanger sequencing at the UC Berkeley DNA Sequencing Facility (Berkeley, CA).

### Comparative gene homolog analyses

Amino acid sequences corresponding to the genes *BdiBd21-3.2G0666400* (*BdPPO1*), *BdiBd21-3.2G0667800* (*BdPPO2*), *BdiBd21-3.2G0668300* (*BdPPO3*), *BdiBd21-3.2G0653800* (*BdTyDC1*), and *BdiBd21-3.2G0654700* (*BdTyDC2*) in *B. distachyon* Bd21-3 v1.1 (https://phytozome-next.jgi.doe.gov/info/BdistachyonBd21_3_v1_1) were searched using DIAMOND (v2.0.152) (Buchfink et al., 2021), with default parameters to identify homologous sequences in the protein databases of Mesangiospermae genomes available on Phytozome (Goodstein et al., 2012). Additionally, proteomes of *Papaver somniferum*, *Musa acuminata*, and *Persea americana* were downloaded from NCBI (accession numbers GCF_003573695.1, GCF_000313855.2, and GCA_029852735, respectively) and included in the search. For species with multiple representatives on Phytozome, such as different cultivars or genome versions, the number of distinct hits for each query protein was averaged across all genome representatives. Homologous hits were considered to be colocalized in the same chromosomal region if the genomic coordinates of at least one pair of genes corresponding to hits from the PPO and TyDC families were located within 10 Mb on the same chromosome. Plant species known to produce dopamine were indicated in Fig. 5A (Gomes et al., 2024; Liu et al., 2020; Novak et al., 2024).

### Sequence analysis and phylogenetic tree construction

Sequence analysis and phylogenetic tree construction were conducted. Protein sequence alignments were performed using Clustal W as implemented in the BioEdit software package (http://www.mbio.ncsu.edu/BioEdit/bioedit.html). Maximum-likelihood phylogenetic trees were constructed using MEGA7 (http://www.megasoftware.net/megabeta.php) with bootstrap values calculated from 1,000 iterations.

## Statistical analysis

Statistical analyses were conducted using JMP Pro v.15.0 (SAS Institute) and Prism v.10.2.3 (GraphPad). All data presented are based on biological replicates, derived from different individual plants. No technical replicates were used for statistical analyses. For two-group comparisons, Student’s unpaired two-tailed *t*-tests were conducted for pairwise comparisons. For comparisons involving more than two groups, one-way ANOVA was performed to evaluate statistical differences. Tukey tests were used to correct for multiple comparisons. For all analyses, *P* < 0.05 was considered to be statistically significant.

## Data and materials availability

Bulk RNA sequence data that support the findings of this study (Supplementary Tables 2, 3, and 4) have been deposited in NCBI Gene Expression Omnibus (GEO; http://www.ncbi.nlm.nih.gov/geo/) with the accession code “GSE284170”. snRNA-seq data are available at NCBI BioProject PRJNA1262374. SNP marker data for mGWAS are available upon request. Processed metabolomics data are provided in Data S1, with references to the corresponding figure panels. Data S2 includes metabolite identifications and GNPS2 links to the raw metabolomics datasets. The Tukey’s multiple-comparison *P* values from one-way ANOVA tests are provided in Data S3. Correspondence and requests for materials should be addressed to T.R.N., J.P.V., or Y.D.

## Author contributions

Y.D. designed the study with input from J.P.V. and T.R.N. Y.D. performed the mapping and functional validation of dopamine biosynthetic genes, with assistance from Y.Z. S.J.L. and S.P.H. performed sequencing for a subset of the *B. distachyon* diversity panel, and L.L. conducted SNP calling for the diversity panel. Metabolomics samples were collected by Y.D. with support from V.N., Y.Z., and S.M.K., and analyzed by Y.D. and S.M.K. Y.D., Y.Z., and M.S. generated CRISPR/Cas9 mutants. Y.D. collected and analyzed the bulk RNA-seq data. T.B. and Y.D. conducted comparative gene homolog analysis. Single-nucleus RNA-seq experiments were performed by L.A.B. and P.W., and analyzed by S.G. Y.D. wrote the manuscript with input from all authors. All authors read and approved the final version of the manuscript.

## Figure legends

**Supplementary Fig. 1.**
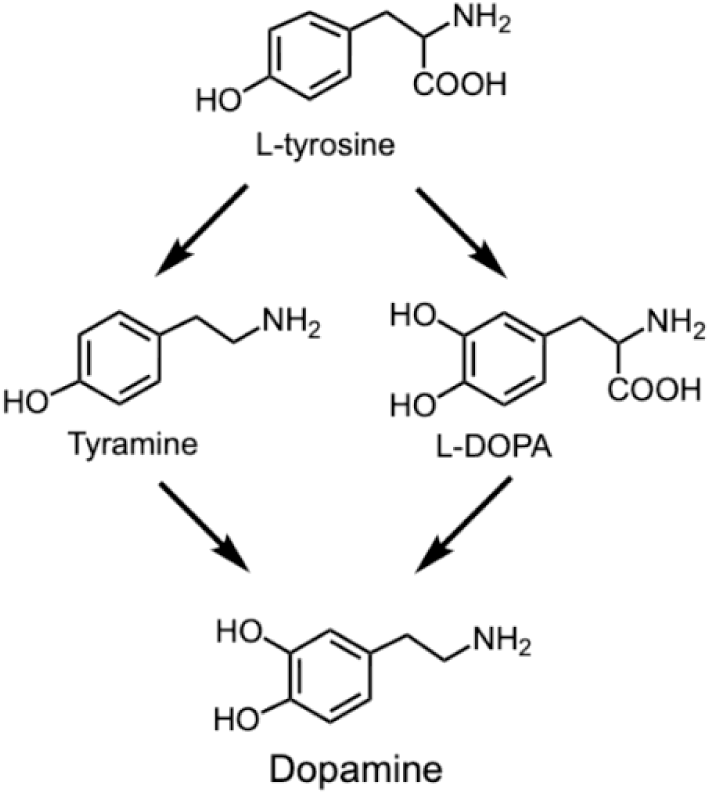
Proposed dopamine biosynthetic pathways in plants.

**Supplementary Fig. 2.**
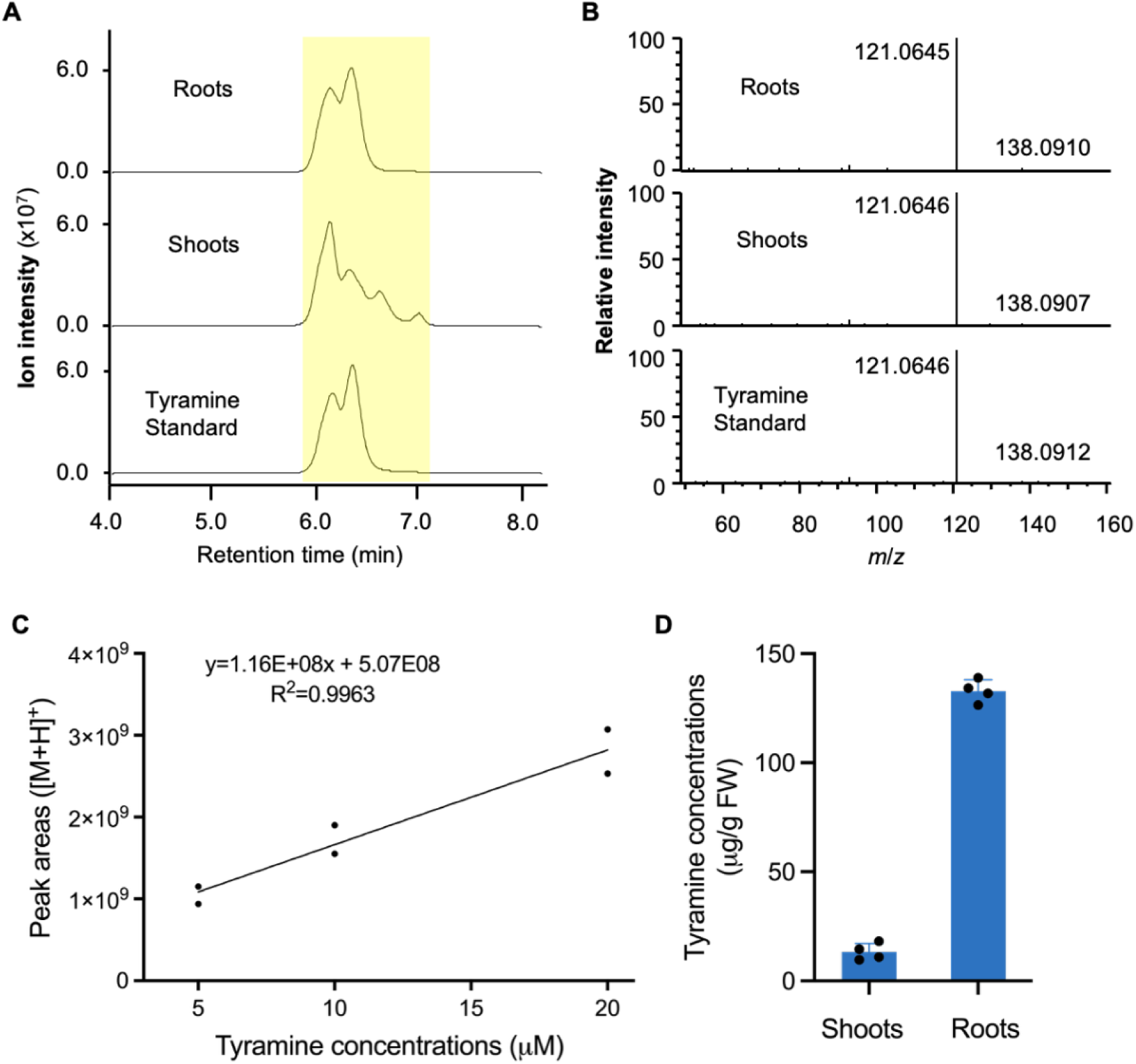
Identification and quantification of tyramine in *B. distachyon* roots and shoots using LC-MS/MS. **(A)** Representative extracted ion chromatograms of tyramine ([M+H]^+^) from *B. distachyon* Bd21-3 roots and shoots, alongside a tyramine standard. **(B)** MS/MS spectra of tyramine detected in roots and shoots compared with the standard. **(C)** External calibration curve for tyramine, generated by plotting the peak area of the standard against its concentration (*n* = 2). **(D)** Quantification of endogenous tyramine levels in Bd21-3 roots and shoots based on the calibration curve shown in (**C**). FW, fresh weight. Error bars represent the mean ± SD (*n* = 4 independent samples).

**Supplementary Fig. 3.**
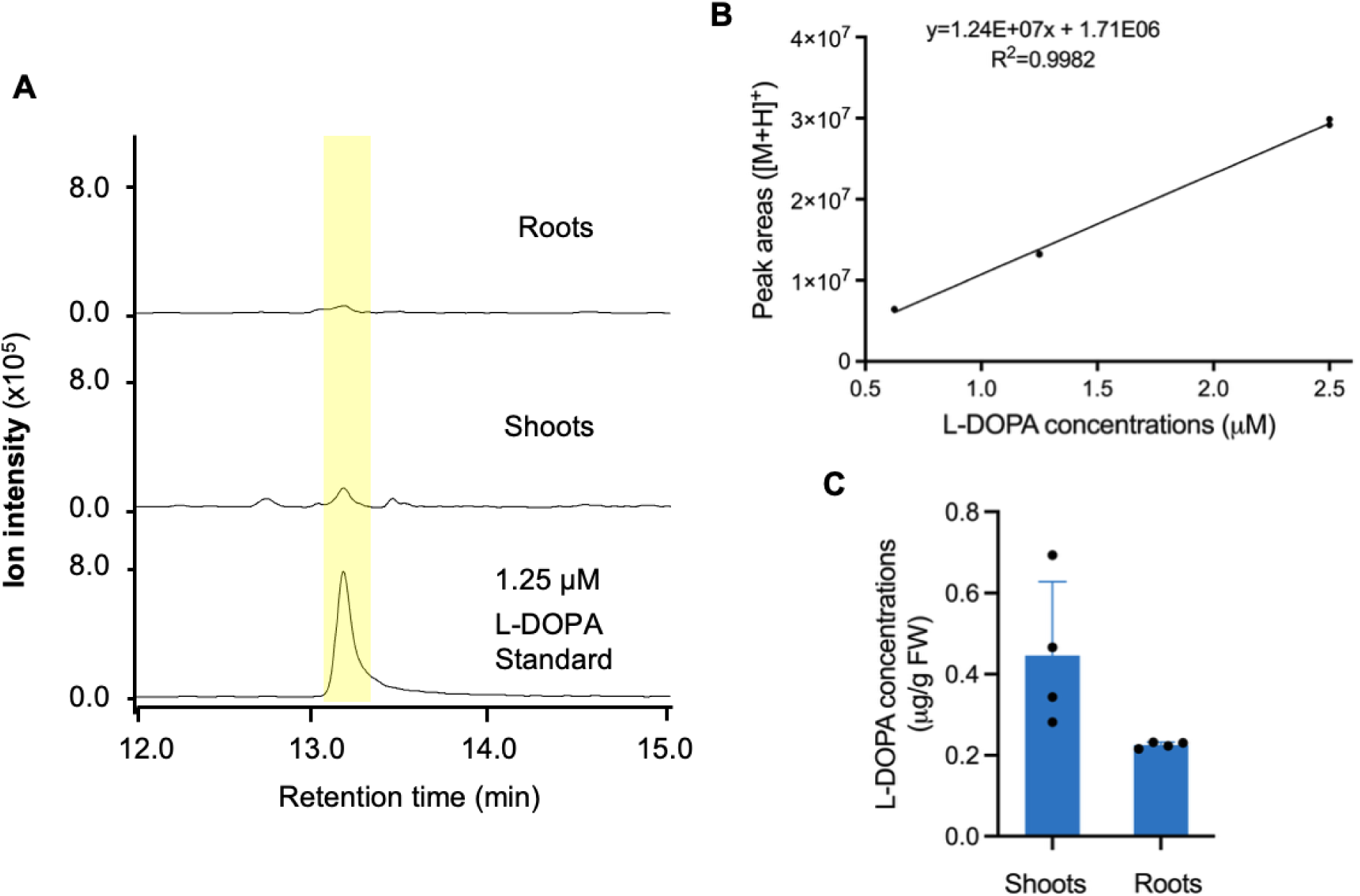
Quantification of L-DOPA in *B. distachyon* roots and shoots using LC-MS/MS. **(A)** Representative extracted ion chromatograms of putative L-DOPA ([M+H]⁺) from *B. distachyon* Bd21-3 roots and shoots, alongside a L-DOPA standard. Due to the presence of only trace amounts, MS/MS spectra could not be acquired for L-DOPA in either tissue. **(B)** External calibration curve for L-DOPA, generated by plotting the peak area of the standard against its concentration (*n* = 2). **(C)** Quantification of endogenous L-DOPA levels in Bd21-3 roots and shoots based on the calibration curve in (**B**). FW, fresh weight. Error bars represent the mean ± SD (*n* = 4 independent samples).

**Supplementary Fig. 4.**
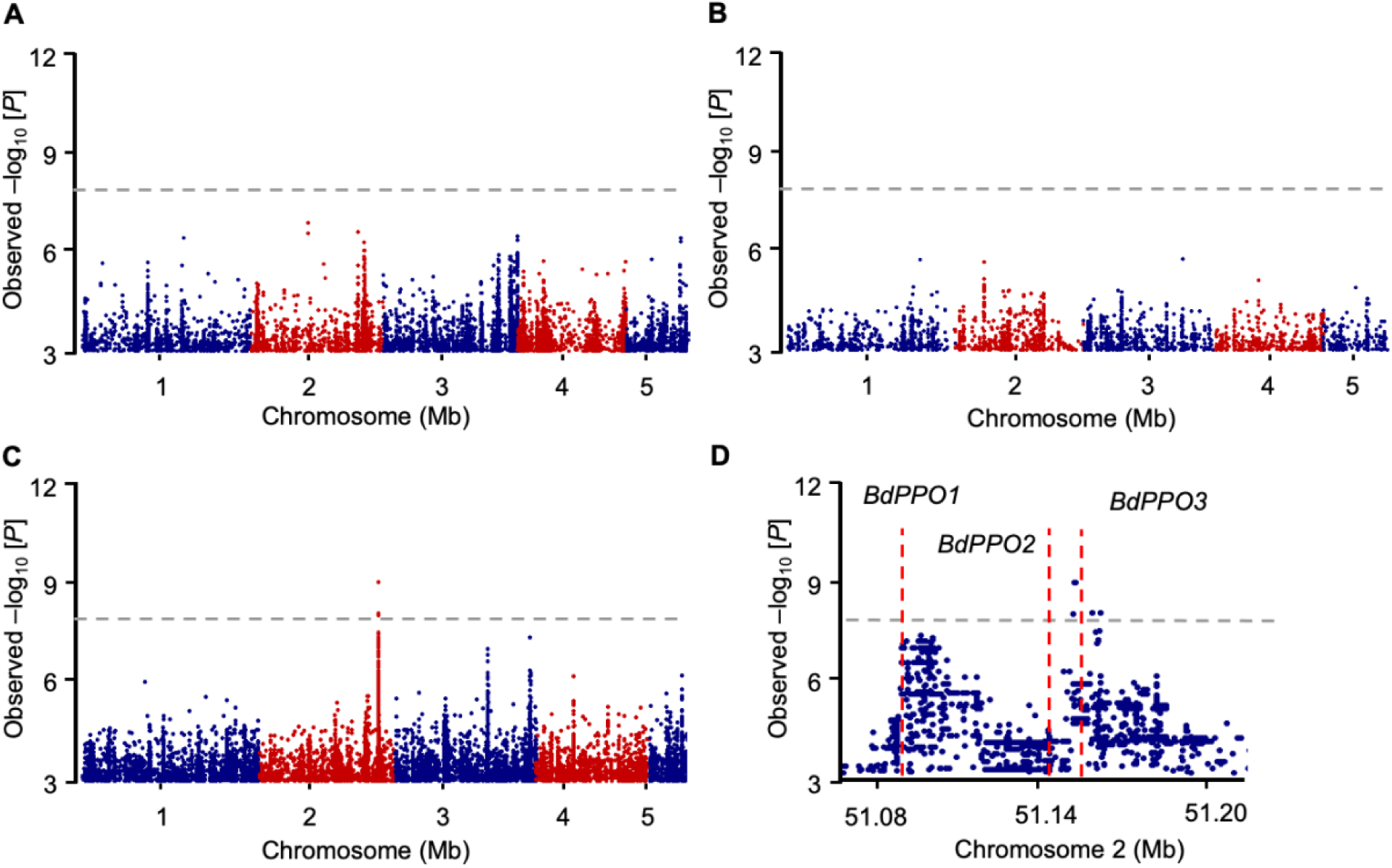
Metabolite-based genome-wide association studies. **(A-C)** Manhattan plots showing metabolite-based GWASs using dopamine levels in roots (**A**) and shoots (**B**) and the dopamine-to-tyramine in shoots (**C**) as mapping traits. Negative log10-transformed *P* values from the general linear model are plotted on the y-axis with the dashed line indicating the 5% Bonferroni-corrected significance threshold (4,780,737 SNPs). The most statistically significant SNPs were identified on chromosome 2. **(D)** A regional Manhattan plot representing a ‘zoomed-in’ view of the signal between 51.05 Mb and 51.25 Mb (Bd21-3 v1.1), and each dot representing a single SNP. The mapping interval overlaps with the region identified for the dopamine-to-tyramine ratio in roots (Fig. 1). Three putative *BdPPO* genes indicated by red dashed lines are located inside the mapping interval with *BdPPO1* (*BdiBd21-3.2G0666400*) in the first peak, *BdPPO3* (*BdiBd21-3.2G0668300*) in the second, and *BdPPO2* (*BdiBd21-3.2G0667800*) between the two peaks. These findings suggest that *BdPPO1* and *BdPPO3* are likely involved in shoot dopamine production.

**Supplementary Fig. 5.**
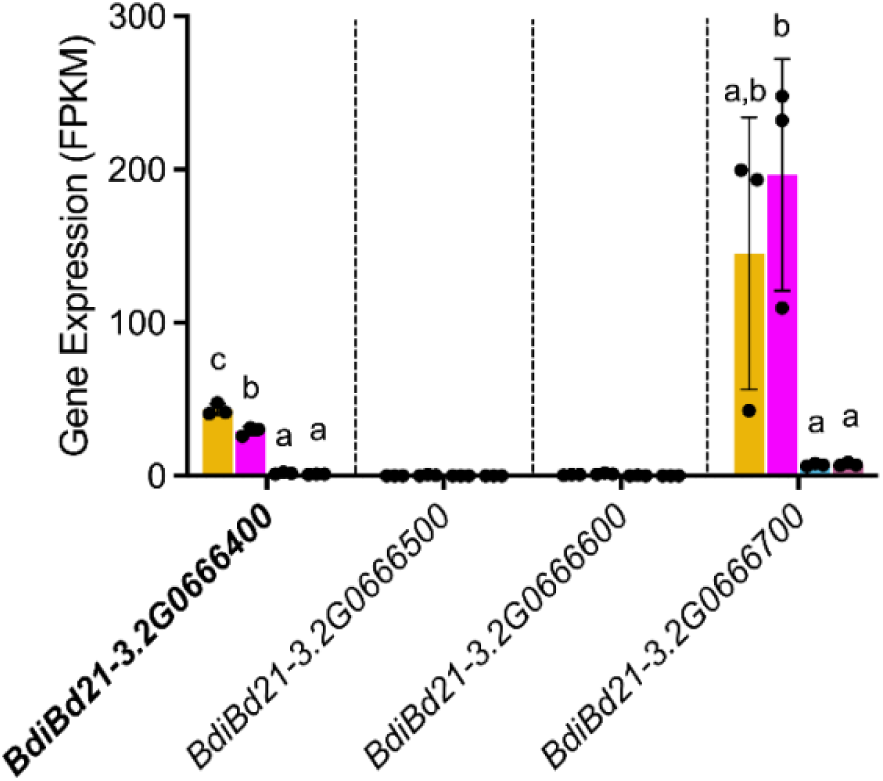
Expression of candidate genes within the genomic interval associated with the root dopamine-to-tyramine ratio. Transcript abundance of four genes (*BdiBd21-3.2G0666400*, *BdiBd21-3.2G0666500*, *BdiBd21-3.2G0666600*, and *BdiBd21-3.2G0666700*) was analyzed by RNA-seq in roots (Root_2wk and Root_4wk) and shoots (Shoot_2wk and Shoot_4wk) of 2-week-old and 4-week-old *B. distachyon* Bd21-3 plants grown hydroponically. Expression values are reported as fragments per kilobase of transcript per million mapped reads (FPKM). Error bars represent mean ± SD (*n* = 3 independent samples). *BdiBd21-3.2G0666500* and *BdiBd21-3.2G0666600* were not detected in any tissue. Among the two expressed genes, *BdiBd21-3.2G0666400*, which encodes a putative polyphenol oxidase, is implicated in dopamine biosynthesis. For each gene, different letters (a-c) indicate statistically significant differences between samples (one-way ANOVA followed by Tukey’s post hoc test; *P* < 0.05).

**Supplementary Fig. 6.**
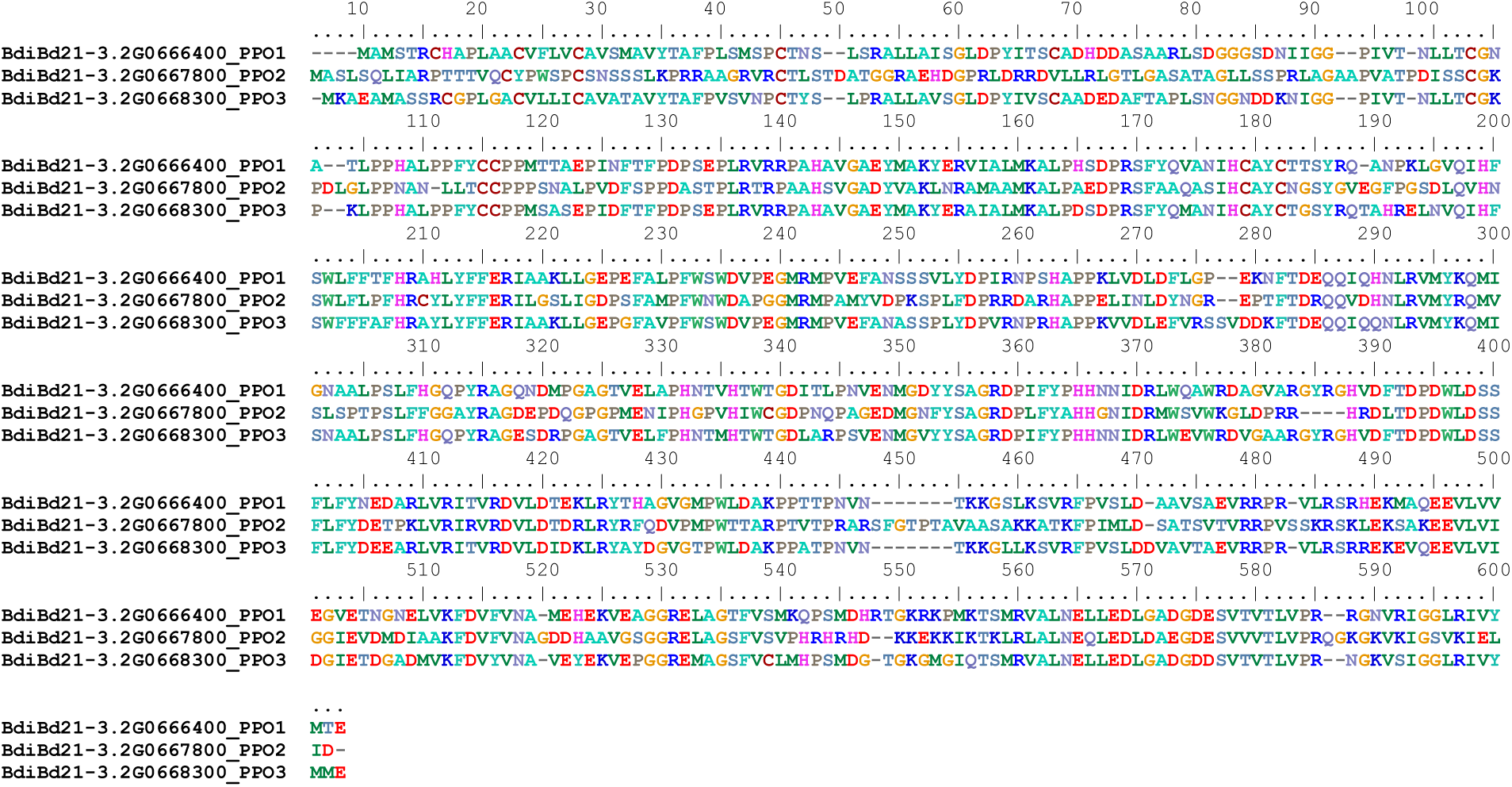
Amino acid sequence comparison of three *B. distachyon* PPOs. The predicted amino acid sequences of three putative *B. distachyon* polyphenol oxidases (PPOs) were aligned using the MUSCLE (codon) algorithm in MEGA7 (www.megasoftware.net). The alignment was visualized with BIOEDIT (http://www.mbio.ncsu.edu/BioEdit). Sequence identity analysis revealed that BdPPO1 shares 52.1% and 78.9% protein sequence identity with BdPPO2 and BdPPO3, respectively, while BdPPO2 and BdPPO3 exhibit 51.0% identity to each other.

**Supplementary Fig. 7.**
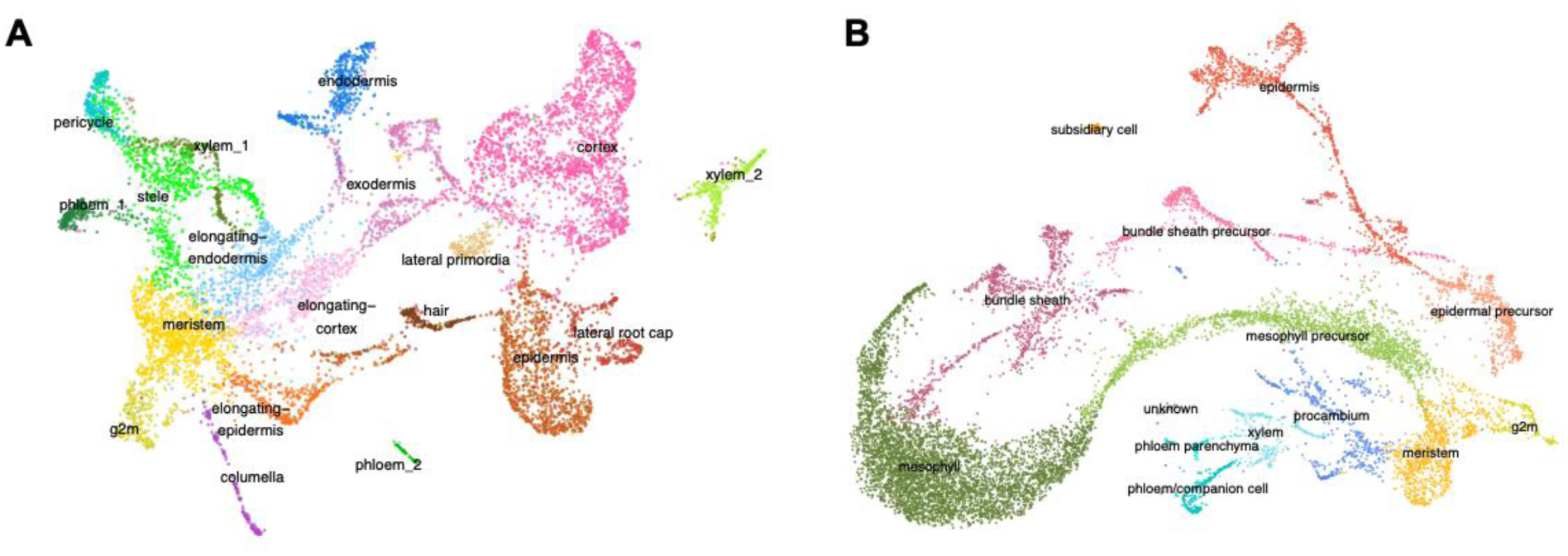
UMAP plots showing cell types in *B. distachyon* roots and shoots. **(A)** UMAP of 9,700 single nuclei from 8-day-old *B. distachyon* roots. **(B)** UMAP of 12,234 single nuclei from 8-day-old *B. distachyon* shoots. Cell types were annotated based on published marker gene sets from maize, *Sorghum bicolor*, and rice.

**Supplementary Fig. 8.**
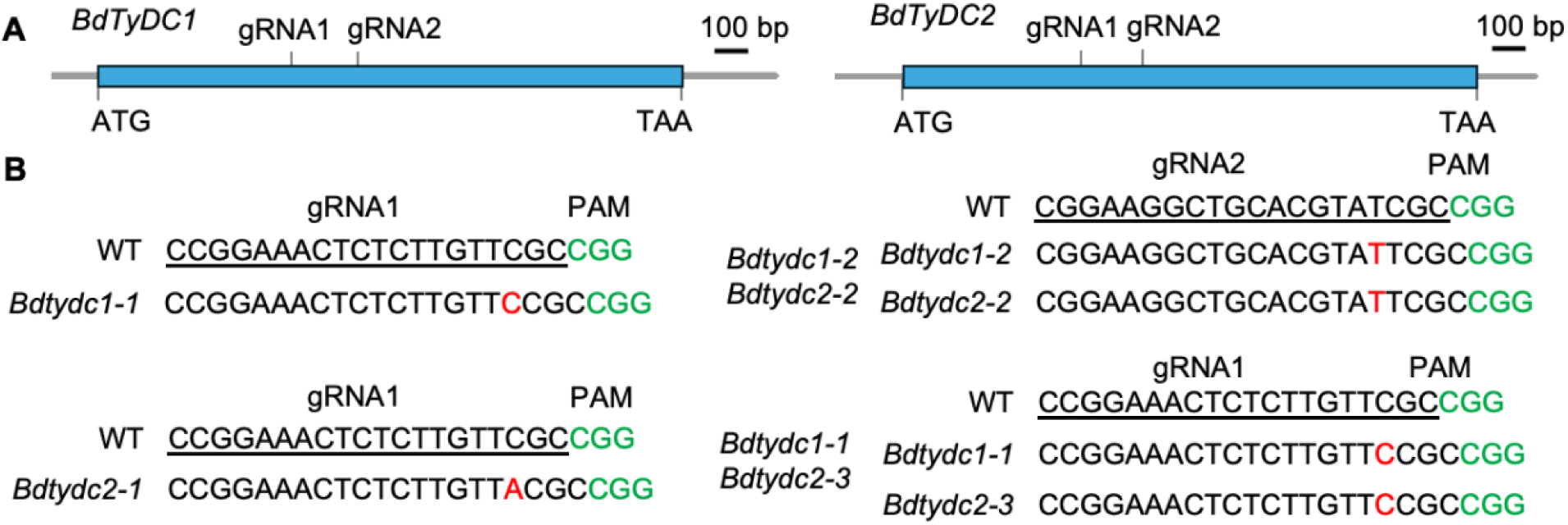
CRISPR/Cas9-induced mutations in *BdTyDC1* and *BdTyDC2.* **(A)** Gene structures of *BdTyDC1* (*BdiBd21-3.2G0653800*) and *BdTyDC2* (*BdiBd21-3.2G0654700*) showing the positions of two guide RNAs (gRNA1 and gRNA2) designed to target both *BdTyDC* genes in Bd21-3. **(B)** Mutations induced by CRISPR/Cas9 at the two *BdTyDC* gene loci. The gRNA target sites are underlined in black, Protospacer adjacent motifs (PAMs) are highlighted in green, and red-highlighted nucleotides indicate insertions.

**Supplementary Fig. 9.**
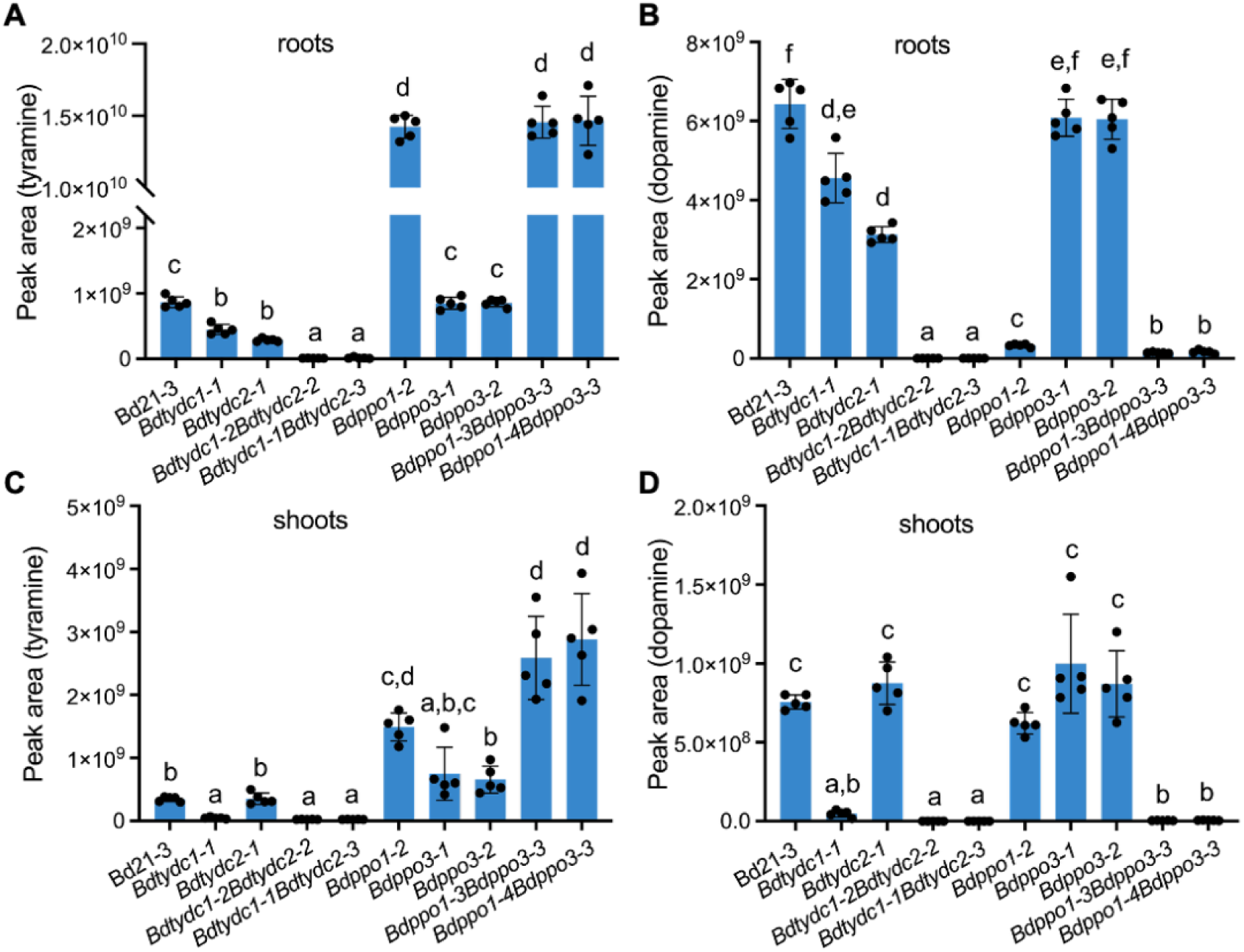
Characterization of dopamine pathway gene mutants. LC-MS/MS peak areas of tyramine ([M+H]⁺) and dopamine ([M+H]⁺) in the roots (**A** and **B**) and shoots (**C** and **D**) of wild-type *B. distachyon* Bd21-3 (wildtype) and various dopamine pathway mutants, including *Bdtydc1-1*, *Bdtydc2-1*, two alleles of *Bdtydc1Bdtydc2*, *Bdppo1-2*, two alleles of *Bdppo3*, and two alleles of *Bdppo1Bdppo3.* Plants were grown hydroponically for 4 weeks prior to tissue harvest and LC-MS/MS analysis. Error bars indicate mean ± SD (*n* = 5 independent samples). Within the plot, different letters (a-d) represent significant differences (one-way ANOVA followed by Tukey’s post hoc test; *P* < 0.05).

**Supplementary Fig. 10.**
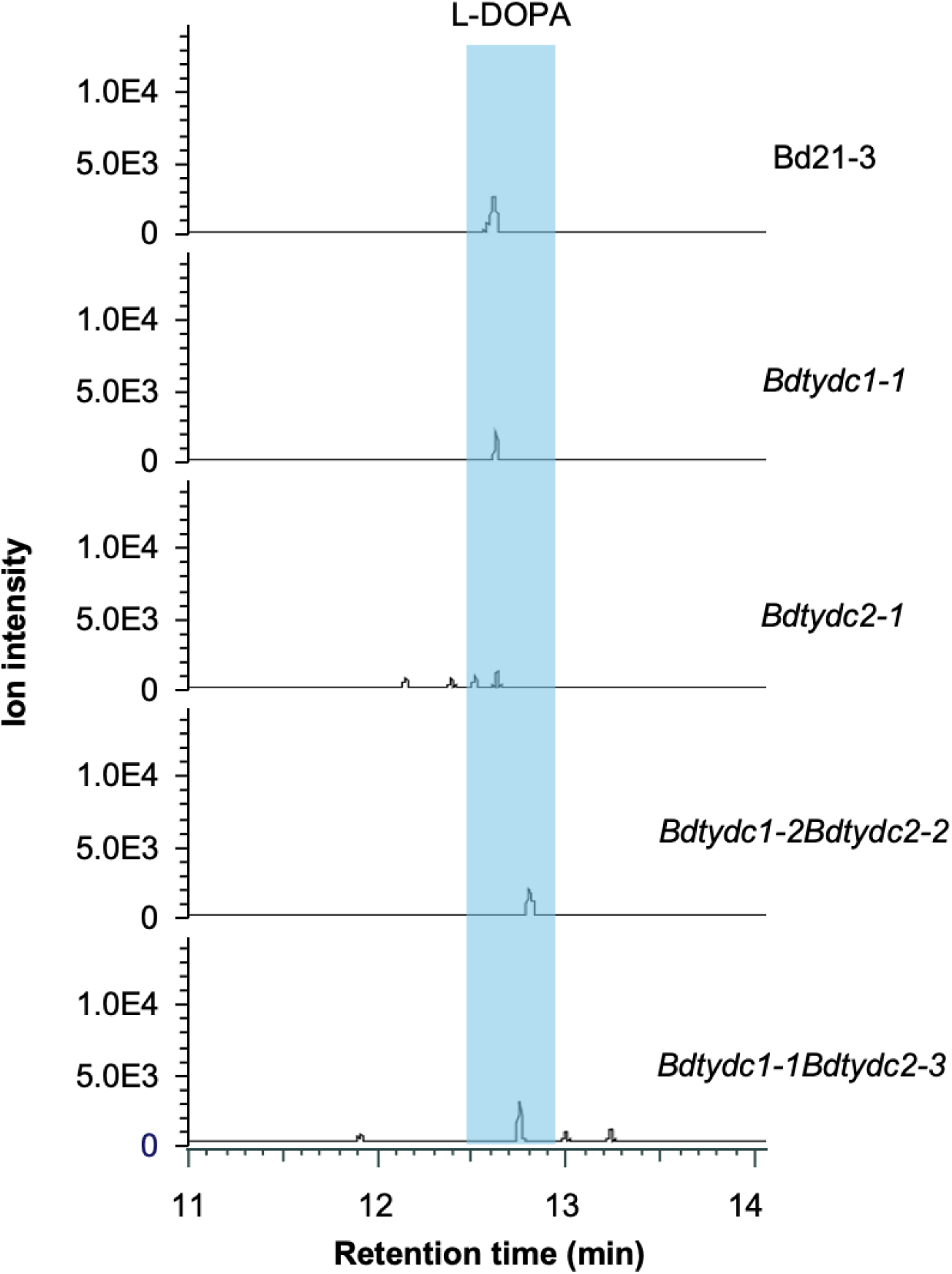
Trace levels of L-DOPA in roots of *Bdtydc* single and double mutants. Representative LC-MS extracted ion chromatograms of L-DOPA ([M+H]⁺) in roots of Bd21-3 (wildtype) and *Bdtydc* mutants, including *Bdtydc1-1*, *Bdtydc2-1*, and two independent alleles of the *Bdtydc1Bdtydc2* double mutant. Plants were grown hydroponically for 4 weeks, and roots were harvested for metabolite analysis via LC-MS/MS. A low but detectable level of putative L-DOPA was consistently observed in roots of all genotypes. Notably, *Bdtydc* single and double mutants did not accumulate more L-DOPA in roots compared to Bd21-3 (wildtype) plants.

**Supplementary Fig. 11.**
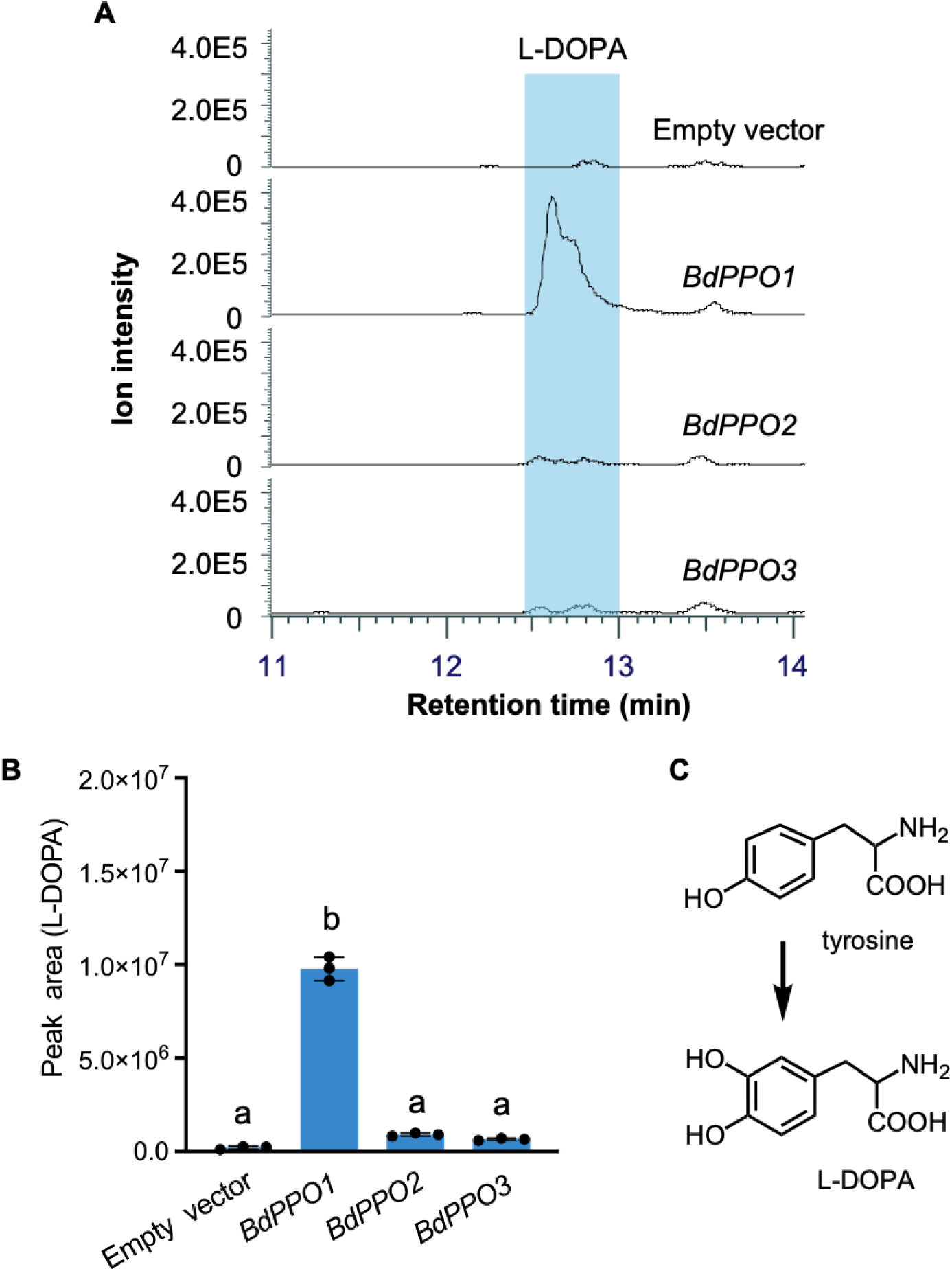
BdPPO1 exhibits minor activity for the 3-hydroxylation of tyrosine. **(A)** Representative LC-MS/MS extracted ion chromatograms of L-DOPA ([M+H]⁺) from *N. benthamiana* leaves transiently expressing *BdPPO1*, *BdPPO2*, *BdPPO3*, or an empty vector. **(B)** LC-MS/MS peak areas of L-DOPA in *N. benthamiana* leaves. Data represent mean ± SD of three biological replicates (*n* = 3 independent samples). A low but detectable level of putative L-DOPA was observed in *BdPPO1*-expressing leaves, whereas BdPPO2 and BdPPO3 did not produce significantly different levels compared to the empty vector control. Different letters (a-b) indicate statistically significant differences (one-way ANOVA followed by Tukey’s post hoc test; *P* < 0.05). **(C)** Proposed enzymatic reaction catalyzed by BdPPO1 in the conversion of tyrosine to L-DOPA.

**Supplementary Fig. 12.**
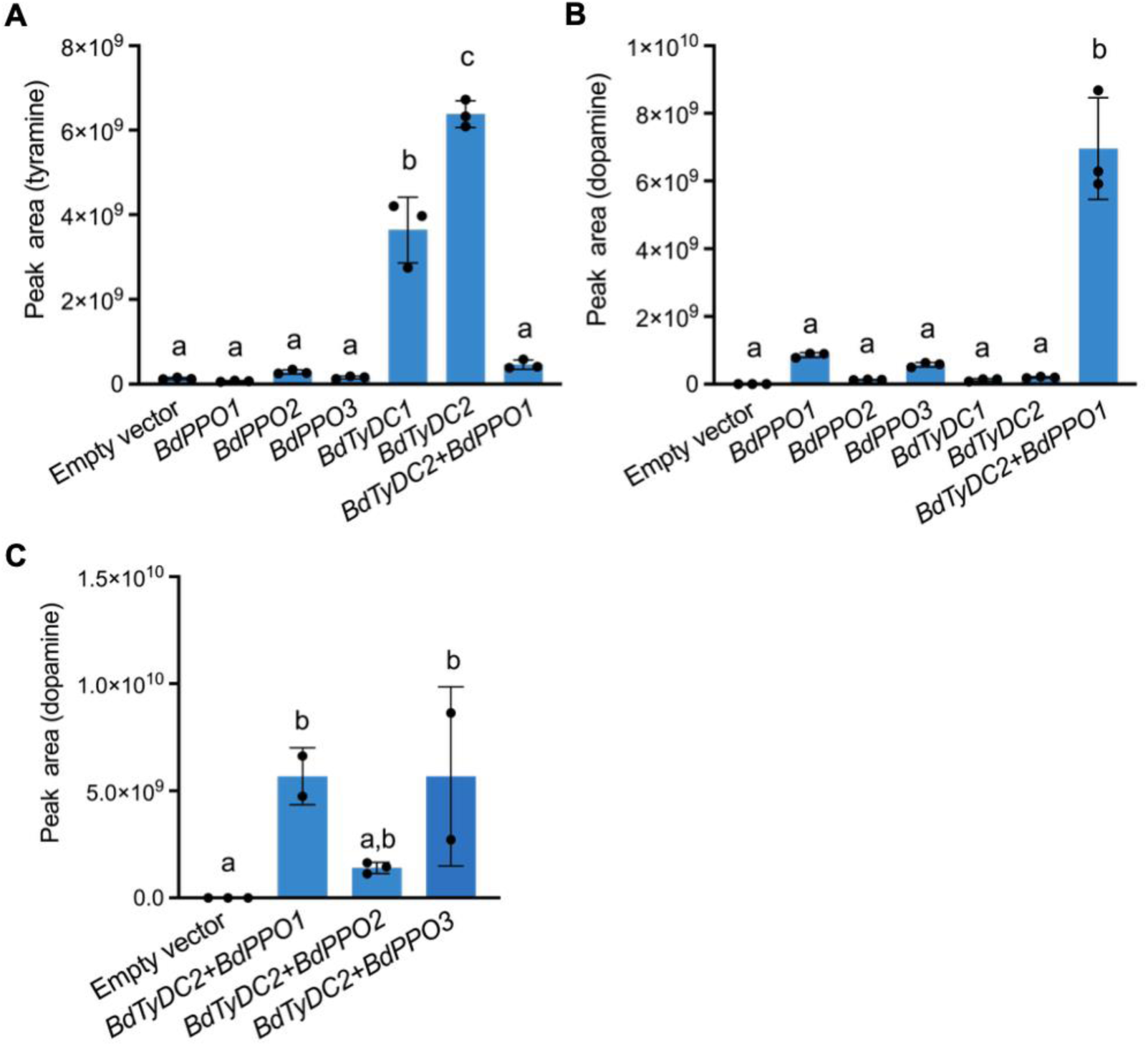
Dopamine and tyramine production in *N. benthamiana* leaves transiently expressing dopamine biosynthetic genes. (**A-B**) LC-MS/MS peak areas of tyramine ([M+H]^+^) (**A**) and dopamine ([M+H]⁺) (**B**) in *N. benthamiana* leaves transiently expressing an empty vector, *BdPPO1*, *BdPPO2*, *BdPPO3*, *BdTyDC1*, *BdTyDC2*, or a combination of *BdTyDC2* and *BdPPO1*. **(C)** LC-MS/MS peak areas of dopamine in leaves co-expressing *BdTyDC2* with *BdPPO1*, *BdPPO2*, or *BdPPO3*. Error bars indicate mean ± SD (*n* = 2 or 3 independent samples). Different letters (a-c) indicate statistically significant differences among groups (one-way ANOVA followed by Tukey’s post hoc test; *P* < 0.05).

**Supplementary Fig. 13.**
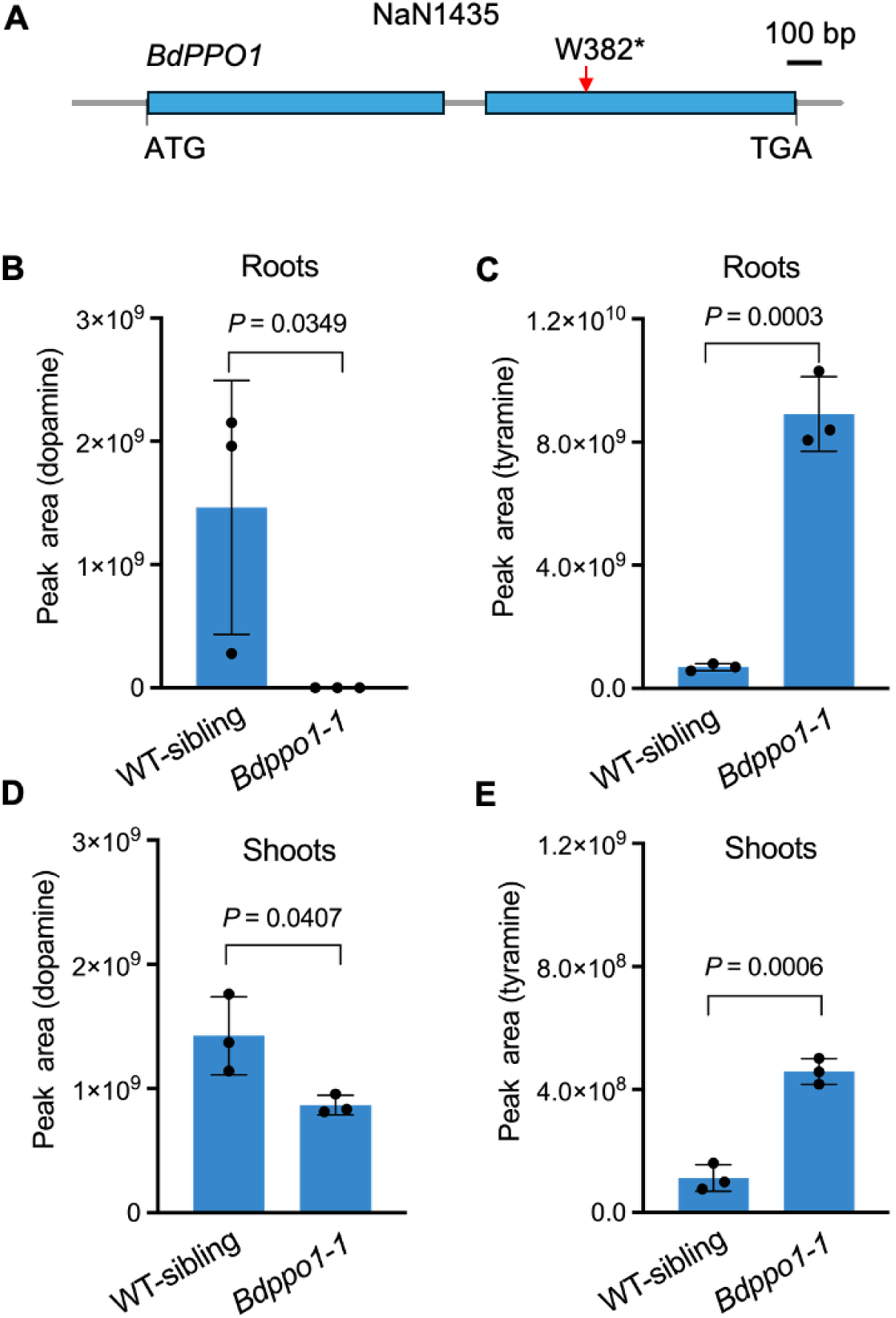
Dopamine and tyramine levels in *Bdppo1-1*. **(A)** Gene structure of *BdPPO1* (*BdiBd21-3.2G0666400*). The NaN1435 mutant line contains a sodium azide-induced mutation in *BdPPO1*, resulting in a premature stop codon (*) at tryptophan 382 (W382). (**B-E**) Dopamine ([M+H]^+^) and tyramine levels ([M+H]^+^) in the roots (**B** and **C**) and shoots (**D** and **E**) of wildtype sibling (WT-Sibling), and *Bdppo1-1.* Plants were grown in a hydroponic system for 4 weeks, and roots and shoots were harvested for untargeted metabolite analysis using LC-MS/MS. Error bars represent mean ± SD (*n* = 3 independent samples). Statistical analysis was performed using Student’s *t*-test.

**Supplementary Fig. 14.**
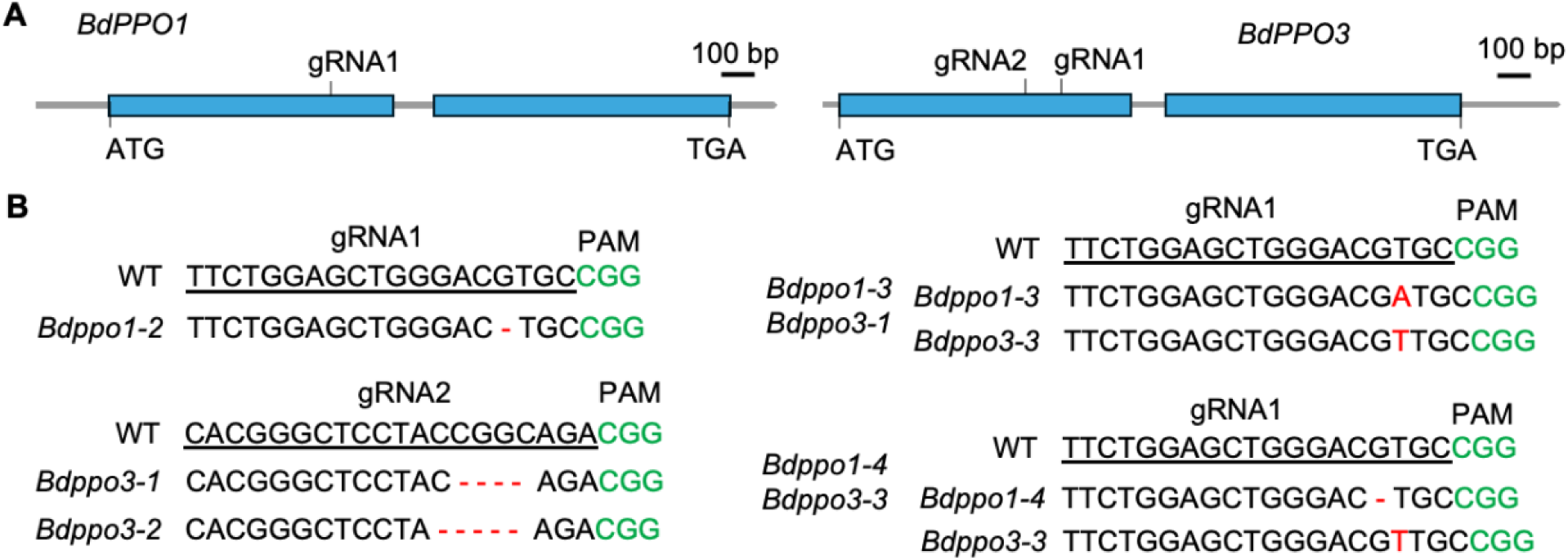
CRISPR/Cas9-induced mutations in *BdPPO1* and *BdPPO3*. **(A)** Schematic representation of *BdPPO1* (*BdiBd21-3.2G0666400*) and *BdPPO3* (*BdiBd21-3.2G0668300*), with locations of two guide RNAs. gRNA1 was designed to target both genes in Bd21-3, while gRNA2 was designed to only target *BdPPO3*. **(B)** Mutations induced by CRISPR/Cas9 at the two gene loci, with gRNA target sites underlined in black. PAMs are marked in green. Red-highlighted nucleotides indicate insertions, and red dashes (– represent nucleotide deletions.

**Supplementary Fig. 15.**
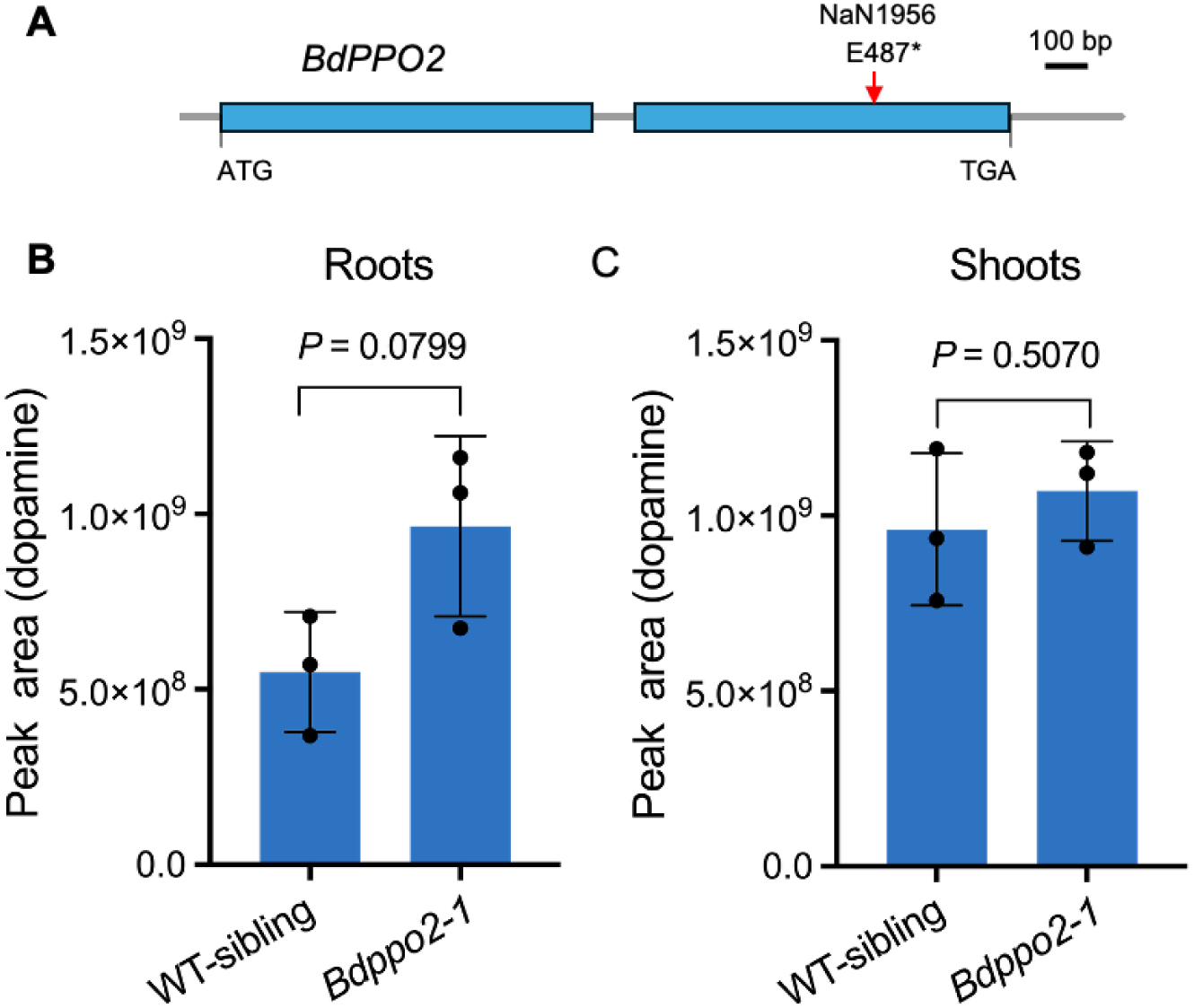
Dopamine levels in *Bdppo2*. **(A)** Gene structure of *BdPPO2* (*BdiBd21-3.2G0667800*). The NaN1956 mutant line contains a sodium azide-induced mutation in *BdPPO2*, resulting in a premature stop codon (*) at glutamic acid 487 (E487). **(B-C)**, Dopamine levels ([M+H]^+^) in the roots (**B**) and shoots (**C**) of the *Bdppo2-1* mutant and its corresponding wildtype sibling (WT-sibling). Plants were grown in a hydroponic system for 4 weeks, and roots and shoots were harvested for LC-MS/MS analysis. Error bars represent mean ± SD (*n* = 3 independent samples). Statistical analysis was performed using Student’s *t*-test.

**Supplementary Fig. 16.**
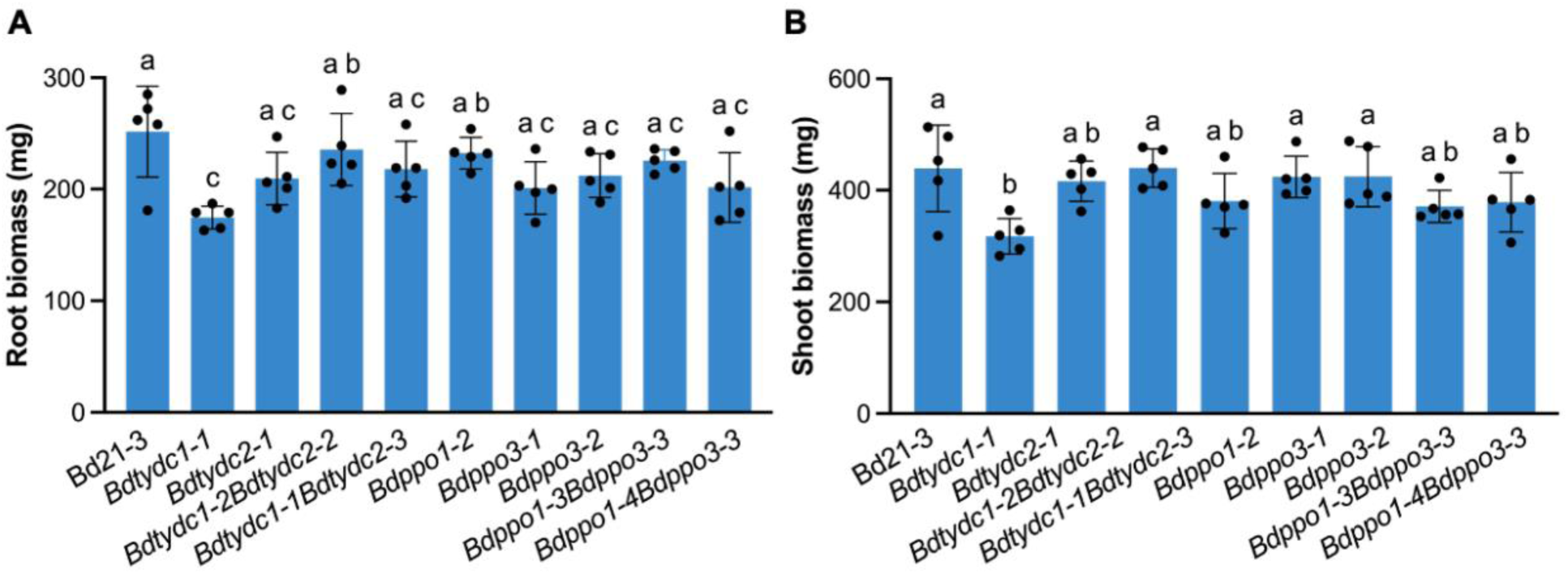
Root and shoot biomass of dopamine pathway mutants. Root (**A**) and shoot (**B**) biomass were measured at week 4. Error bars represent mean ± SD (*n* = 5 independent samples). Plants, including Bd21-3 (wildtype), *Bdtydc1-1*, *Bdtydc2-1*, two alleles of *Bdtydc1Bdtydc2*, *Bdppo1-2*, two alleles of *Bdppo3*, and two alleles of *Bdppo1Bdppo3* were grown in a hydroponic system. Within the plot, different letters (a-c) represent significant differences (one-way ANOVA followed by Tukey’s test corrections for multiple comparisons; *P* < 0.05).

**Supplementary Fig. 17.**
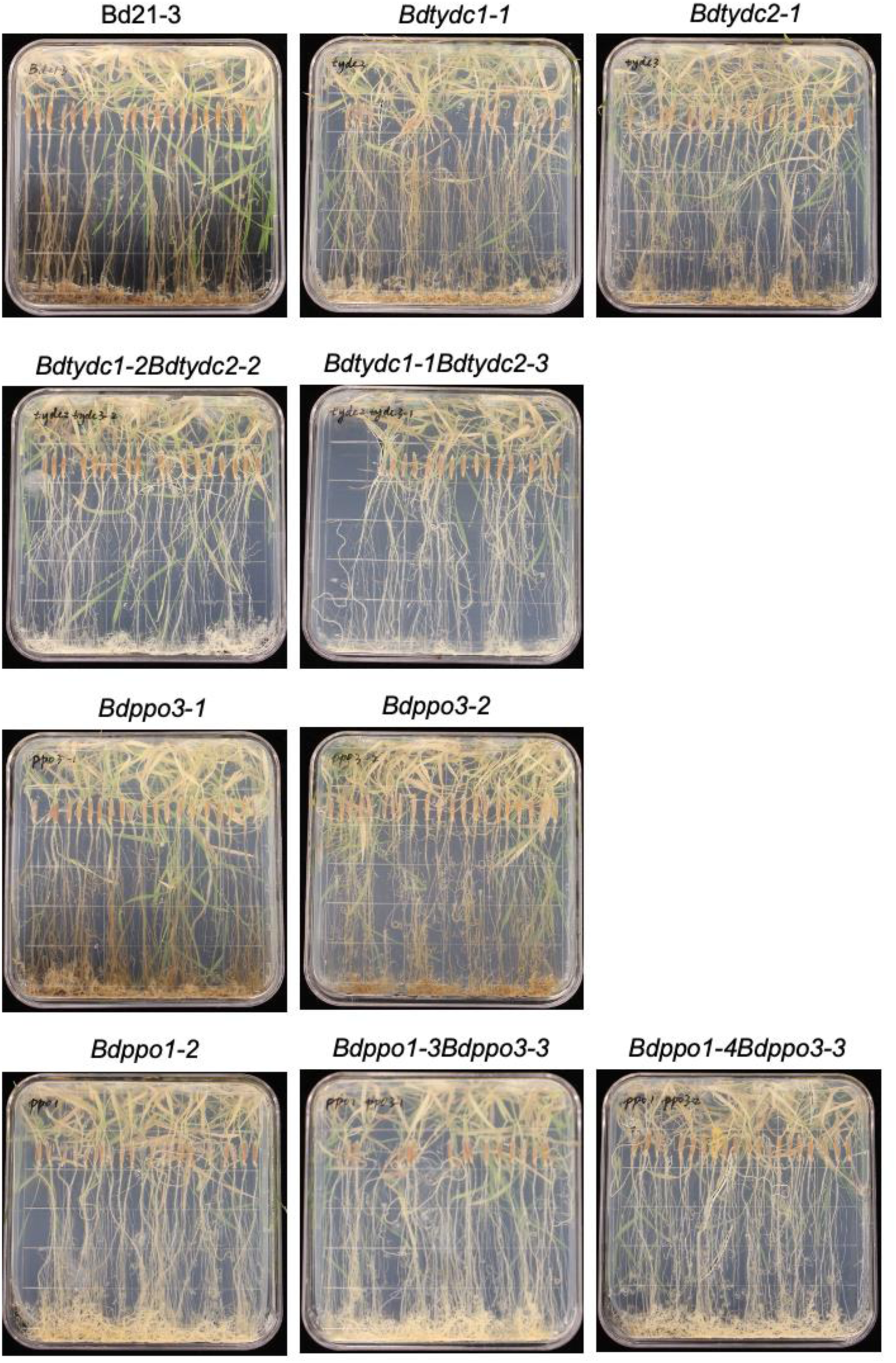
Senescence-induced root browning in *B. distachyon*. Plants, including Bd21-3 (wildtype) and dopamine pathway mutants (*Bdtydc1-1*, *Bdtydc2-1*, two alleles of *Bdtydc1Bdtydc2*, *Bdppo1-2*, two alleles of *Bdppo3*, and two alleles of *Bdppo1Bdppo3*) were grown on ½ MS plates for approximately 8 weeks before images were taken.

**Supplementary Fig. 18.**
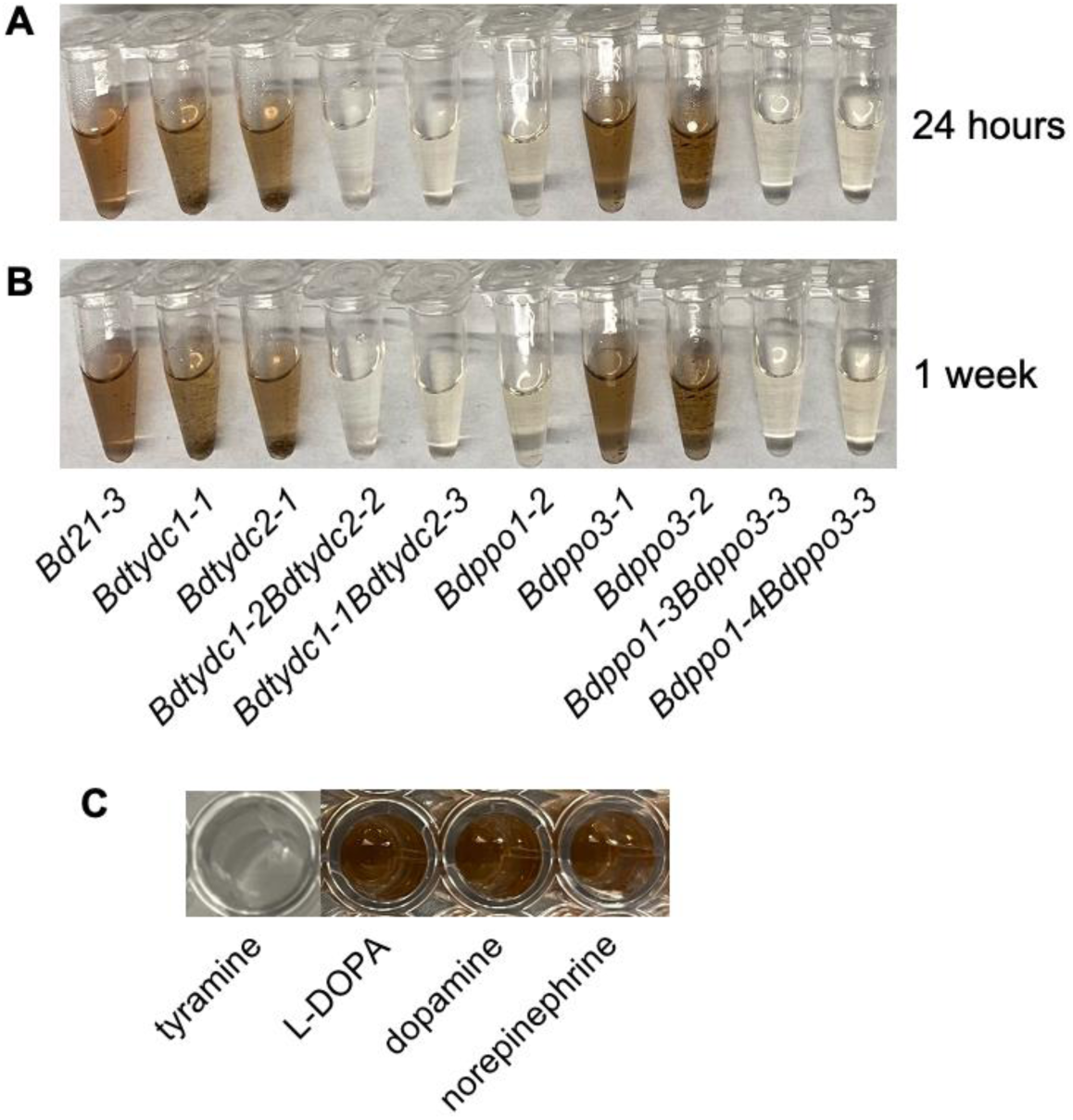
Browning of root extracts. **(A-B)**, Root extracts from three-week-old Bd21-3 (wildtype) and dopamine pathway mutant plants grown on ½ MS plates were exposed to air for 24 hours (**A**) and 1 week (**B**). The dopamine pathway mutants used here include *Bdtydc1-1*, *Bdtydc2-1*, two alleles of *Bdtydc1Bdtydc2*, *Bdppo1-2*, two alleles of *Bdppo3*, and two alleles of *Bdppo1Bdppo3.* **(C)** Autoxidation of dopamine-related compounds. Tyramine, L-DOPA, dopamine, and norepinephrine were each dissolved in water at an approximate concentration of 100 µM and incubated under constant shaking for around 48 hours with exposure to air to monitor autoxidation. L-DOPA and norepinephrine are at trace levels in *B. distachyon* (Ding et al., 2024).

